# Nucleotidyltransferase toxin MenT targets and extends the aminoacyl acceptor ends of serine tRNAs *in vivo* to control *Mycobacterium tuberculosis* growth

**DOI:** 10.1101/2024.03.07.583660

**Authors:** Xibing Xu, Roland Barriot, Bertille Voisin, Tom J. Arrowsmith, Ben Usher, Claude Gutierrez, Xue Han, Peter Redder, Tim R. Blower, Olivier Neyrolles, Pierre Genevaux

## Abstract

Toxins of toxin-antitoxin systems use diverse mechanisms to control bacterial growth and represent attractive therapeutic targets to fight pathogens. In this study, we characterized the translation inhibitor toxin MenT3 of *Mycobacterium tuberculosis*, the bacterium responsible for human tuberculosis in humans. We show that MenT3 is a robust cytidine specific tRNA nucleotidyltransferase *in vitro*, capable of modifying the aminoacyl acceptor ends of most tRNA but with a marked preference for tRNA^Ser^, to which long stretches of cytidines were added. Furthermore, transcriptomic-wide analysis of MenT3 targets in *M. tuberculosis* identified tRNA^Ser^ as the sole target of MenT3 *in vivo* and revealed significant detoxification attempts by ribonuclease PH in response to MenT3 overexpression. Finally, under physiological conditions, only in the presence the native *menAT3* operon, we found the unexpected presence of an active pool of endogenous MenT3 targeting tRNA^Ser^ in *M. tuberculosis*, likely reflecting the importance of MenT3 during infection.

## INTRODUCTION

Toxin-antitoxin (TA) systems are stress responsive genetic elements encoding for a deleterious toxin and its antagonistic antitoxin. They are widespread in bacterial genomes and mobile genetic elements (LeRoux *et al*, 2020; Harms *et al*, 2018; Jurėnas *et al*, 2022). They have roles in defending bacteria against phage infection and in the maintenance of genomic regions and plasmids, and in some cases they have been shown to contribute to bacterial virulence and antibiotic persistence (Pecota & Wood, 1996; Guegler & Laub, 2021; Helaine *et al*, 2014; Dedrick *et al*, 2017; De Bast *et al*, 2008; Fineran *et al*, 2009). In the absence of stress, toxin activity is blocked by its antagonistic antitoxin and bacterial growth is not detectably impacted. Yet, under specific conditions, including phage infection or the loss of plasmids, the toxin and antitoxin equilibrium can be significantly unbalanced in favor of the more stable toxin. As a consequence, the free active toxin can target essential cellular processes or structures, mainly translation, DNA replication, metabolism or the cell envelope, causing growth inhibition or cell death (Jurėnas *et al*, 2022; Harms *et al*, 2018).

Tuberculosis is the second cause of death due to an infectious agent, after COVID-19. According to the most recent WHO report (2023), over 10 million people fell ill with tuberculosis in 2022 and 1.3 million died from the disease. The increasing occurrence of multi and extensively drug-resistant strains of the causative *Mycobacterium tuberculosis* bacterium has greatly heightened the need for the development of new drugs and new treatment strategies. *M. tuberculosis* encodes an unusually high number of TA systems, over 86, representing close to 4% of its genome (Akarsu *et al*, 2019; Sala *et al*, 2014). This includes multiple homologs from conserved TA families that have been shown to be induced under relevant stress conditions, including hypoxia, macrophage engulfment, or drug exposure (Ramage *et al*, 2009; Keren *et al*, 2011; Ariyachaokun *et al*, 2020). Although their contribution to *M. tuberculosis* physiology and virulence is currently unknown, it has been proposed that activated toxins could modulate *M. tuberculosis* growth under certain conditions, thereby contributing to survival in the human host (Keren *et al*, 2011; Barth *et al*, 2021; Sala *et al*, 2014). To date, only a few TA systems have been tested and shown to contribute to host infection (Deep *et al*, 2018; Tiwari *et al*, 2015; Agarwal *et al*, 2018). The deleterious nature of many *M. tuberculosis* toxins has raised the possibility that new antibacterial properties demonstrated by toxins might be exploited to identify new drug targets or applied directly as antimicrobials, alone or in combination with standard antibacterial therapy (Freire *et al*, 2019; Kang *et al*, 2018; Catara *et al*, 2023).

*M. tuberculosis* encodes four MenAT TA systems, comprised of a nucleotidyltransferase (NTase) toxin and a cognate antitoxin belonging to one of three different families (Dy *et al*, 2014; Xu *et al*, 2023; Cai *et al*, 2020). Although, *in vivo*, *menAT2* was recently shown to be required for *M. tuberculosis* pathogenesis in guinea pigs (Gosain *et al*, 2022), only *menAT1* and *menAT3* were shown to act as *bona fide* TA systems in their native host *M. tuberculosis* (Xu *et al*, 2023; Cai *et al*, 2020). The MenA3 antitoxin inhibits MenT3 through phosphorylation of a serine residue in the catalytic site (Yu *et al*, 2020), whilst MenA1 forms an asymmetric heterotrimeric complex with two MenT1 protomers, suggesting a different mode of inhibition (Xu *et al*, 2023). *In vitro*, the MenT1, MenT3 and MenT4 toxins were shown to inhibit translation by acting as tRNA NTases (Xu *et al*, 2023; Cai *et al*, 2020). Yet, biochemical characterization revealed significant differences in MenT toxin specificity for tRNA targets and nucleotide substrates. Though both MenT1 and MenT3 inhibit aminoacylation by transferring pyrimidines (preferentially cytidines) to the 3′ CCA acceptor stems of tRNAs, MenT3 displays a strong preference for serine tRNAs and MenT1 showed no apparent preference (Xu *et al*, 2023; Cai *et al*, 2020). MenT4 also modifies the 3′ CCA motif of tRNA acceptor stems but with a preference for GTP as substrate (Xu *et al*, 2023). We have previously demonstrated that MenT3 is by far the most toxic of the four MenT toxins of *M. tuberculosis*, inducing a rapid and efficient self-poisoning *in vivo* (Cai *et al*, 2020). Despite the accumulated knowledge concerning MenT3 *in vitro* activity and structure, nothing is currently known about MenT3 activity and cellular targets *in vivo* within *M. tuberculosis*.

In this work, we have investigated the impact of MenT3 on *M. tuberculosis*, focusing on the identification of tRNA targets on a transcriptomic scale, on the molecular determinants necessary for tRNA recognition, and on the mechanisms by which *M. tuberculosis* counteracts noxious tRNA modification, both *in vivo* and *in vitro*. Using a combination of tRNA-seq *in vivo* and *in vitro*, and several biochemical approaches, we show that MenT3 is an effective NTase capable of efficiently modifying all the tRNA of *M. tuberculosis in vitro*, by adding 3 to 4 cytidines at the 3′ CCA end of the acceptor stem. Cytidine elongation was much more efficient for tRNA^Ser^, reaching up to 17 cytidines added. Grafting the long variable loop present in all tRNA^Ser^ to unrelated tRNAs promoted the addition of similarly longer stretches of cytidines, indicating that the variable loop is a determinant for optimal MenT3 activity. Remarkably, when expressed *in vivo* in *M. tuberculosis*, MenT3 specifically and efficiently added cytidines to all tRNA^Ser^, without any modification detected for other tRNAs. In sharp contrast with results *in vitro*, we found that different tRNA^Ser^ 3′ end species were accumulating *in vivo*, mostly deleted for the third position adenosine, resulting in CCΔA, CΔCA and CCΔA+C*n* extensions instead of CCA+C*n*. Such a phenomenon was fully recapitulated *in vitro* in the presence of *M. tuberculosis* exoribonuclease RNase PH, which appears to be a key factor in the cellular response to MenT3 activity. Importantly, we could also identify a steady state level of cytidine modification of tRNA^Ser^ in *M. tuberculosis* wild type under standard growth conditions, and thus solely in the presence of the native chromosomal *menAT3* operon copy. The role of such unexpected basal level of MenT3-dependent tRNA^Ser^ modification and its possible link with *M. tuberculosis* physiology and virulence is discussed.

## RESULTS

### MenT3 is a promiscuous tRNA NTase with robust poly-C activity *in vitro*

We previously showed that MenT3 can add two to three cytidines or uridines to *in vitro* transcribed tRNAs (Cai *et al*, 2020). In addition, individual screening of each of the 45 *in vitro* purified tRNA of *M. tuberculosis* showed that MenT3 could modify the four tRNA^Ser^ and to a lesser extent tRNA^Leu-5^, while all the other tRNAs were not detectably modified. Yet, our recent observation that mature tRNA were more efficiently modified *in vitro* by the related MenT1 toxin (Xu *et al*, 2023) led us to reinvestigate the nucleotide and tRNA specificities of MenT3. Incubation of *M. smegmatis* tRNA extract with MenT3 in the presence of each individual labeled nucleotide indicates that although UTP (and to a much lesser extent ATP) can be used by MenT3, CTP is by far the preferred nucleotide substrate of MenT3 *in vitro* (**Fig. 1A**). In addition, we found that MenT3 is capable of efficiently modifying *M. tuberculosis*, *E. coli* and human tRNAs purified from cell extracts, indicating that MenT3 is a promiscuous tRNA NTase *in vitro* (**Fig. 1B**). Furthermore, the NTase activity of MenT3 towards tRNA extracts *in vitro* is significantly more robust than those of MenT1 or MenT4 (**Fig. 1C**), which is in agreement with the strong toxicity of MenT3 *in vivo*, when compared to the other MenT toxins tested so far (Cai *et al*, 2020; Xu *et al*, 2023).

**Fig. 1:**
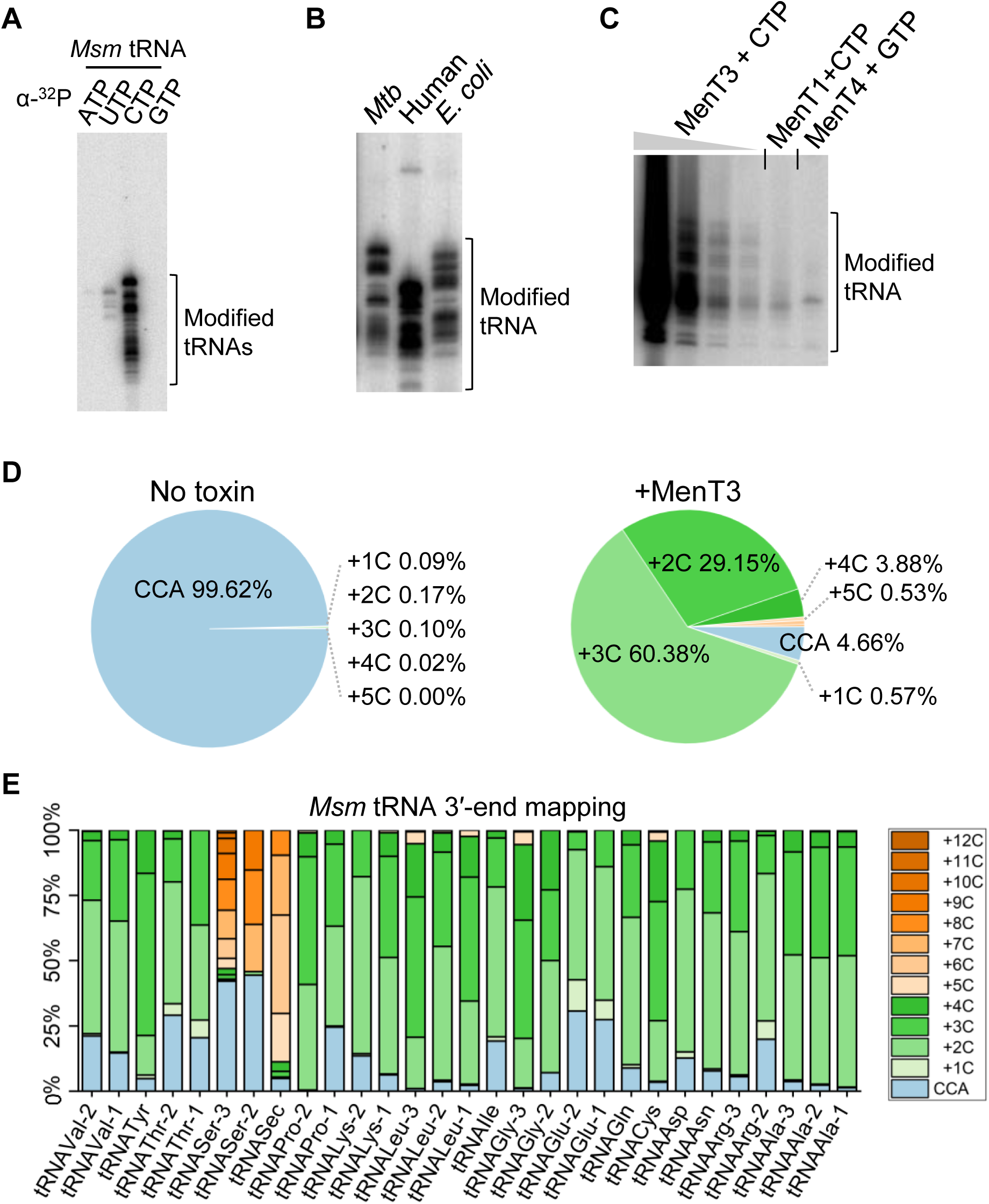
MenT3 NTase activity *in vitro*. (**A**) MenT3 preferentially adds CTP to tRNA. Total tRNA (100 ng) of *M. smegmatis* (*Msm*) were incubated at 37 ℃ for 5 min with MenT3 (0.2 µM) and α-P32 labeled nucleoside triphosphates (NTPs). (**B**) Promiscuous tRNA modification by MenT3. 1 µg of total RNA from *M. tuberculosis* (*Mtb*) or human cells, or 100 ng of tRNA extracts from *E. coli* were incubated with 0.2 µM MenT3 in the presence of α-P32 labeled CTPs for 5 min at 37 ℃. (**C**) Comparison of tRNA NTase activity among MenT toxins. MenT3 (0.2 µM) or MenT1 (5 µM), or MenT4 (5 µM) were incubated with 1 µg of total RNA of *Msm* for 20 min at 37 ℃. Given the robust activity of MenT3, the sample was serial diluted 1/10, 1/100 or 1/1000 to facilitate visualization of MenT1 and MenT4 activity through phosphor exposure. Reactions were conducted in the presence of α-P32 labeled CTP for MenT1 and MenT3, and α-P32 labeled GTP for MenT4. Representative results of triplicate experiments are shown in panels A, B and C. (**D**) and (**E**) tRNA 3′-end mapping. Five µg total RNA from *Msm* were incubated with MenT3 (2.5 µM) and 1 mM CTP for 20 minutes at 37 ℃, and the tRNA-seq library was prepared and sequenced. Modified tRNA reads were quantified in samples with or without MenT3 and the modifications are given as a percentage of the total tRNA 3′-ends. Note that tRNA^Sec^ is not present in *M. tuberculosis*. The tRNA detected from two independent experiments are shown on the bottom of panel E and the number of cytidines added is shown in the inset at the right of the same panel. (Duplicate is shown in datasheet file).

We next applied 3′-OH tRNA-seq to the *M. smegmatis* RNA extracts following incubation in the presence or absence of MenT3 and found that over 95% of the pool of detected tRNA was modified with an average of two to five cytidines added to 3′CCA tRNA ends (CCA+C*n*; **Fig. 1D**). Remarkably, while most of the tRNAs had these two to five cytidine extensions, the detected tRNA^Ser2^ and tRNA^Ser3^ showed a completely different behavior, with significantly longer poly-cytidine (poly-C) extensions of up to *n*=12 under such conditions (**Fig. 1E, Supplementary Datasheet**). Note that tRNA^Ser-1^ and tRNA^Ser-4^ could not be sufficiently detected in *M. smegmatis* extracts. In addition, the *M. smegmatis* selenocysteine tRNA^Sec^, which is not present in *M. tuberculosis* (Behra *et al*, 2022) and is very similar to tRNA^Ser^ was also highly modified by MenT3. Together these *in vitro* data show that MenT3 is a very robust CTP-dependent tRNA NTase toxin, which can modify most tRNAs with 3′CCA+C*_(2-5)_* extensions, but with a preference for tRNA^Ser^ (or tRNA^Sec^ in *M. smegmatis*) with respect to the length of the cytidine extension.

### Serine tRNA variable loop is critical for poly-C extensions *in vitro*

We further investigated the marked difference in cytidine extension observed between tRNA^Ser^ and other tRNAs *in vitro*, using purified *M. tuberculosis* tRNA^Ser-4^ and tRNA^Met-2^ as representative tRNAs. In this case, the tRNAs were independently labelled with [α-^32^P]-CTP and purified using a ribozyme-based cleavage method that generated more homogeneous 3′-OH ends, to avoid the higher heterogeneity generated in transcripts made with T7 RNA polymerase transcription (Xu *et al*, 2023). Labeled tRNAs were individually incubated with MenT3 in the presence of CTP and samples were analyzed at different time points (**Fig. 2A**). The data show the addition of cytidines in both cases, with a more rapid accumulation of longer species of modified tRNA^Ser-4^ when compared to tRNA^Met-2^. Analysis of tRNA^His^ and tRNA^Leu-3^ confirmed the accumulation of shorter extensions as observed for tRNA^Met-2^ (**Supplementary Figure S1**). Furthermore, 3′-OH tRNA-seq analysis of the 10 min incubation samples of tRNA^Ser-4^ and tRNA^Met-2^ revealed that the majority of the tRNA^Ser-4^ have indeed acquired 11 to 12 cytidines (with a maximum n=17), while tRNA^Met-2^ only two to five (with a majority at n=3) (**Fig. 2B**). Together these *in vitro* data demonstrate that although MenT3 is capable of modifying most tRNA *in vitro*, it shows a strong preference for tRNA^Ser^.

**Fig. 2:**
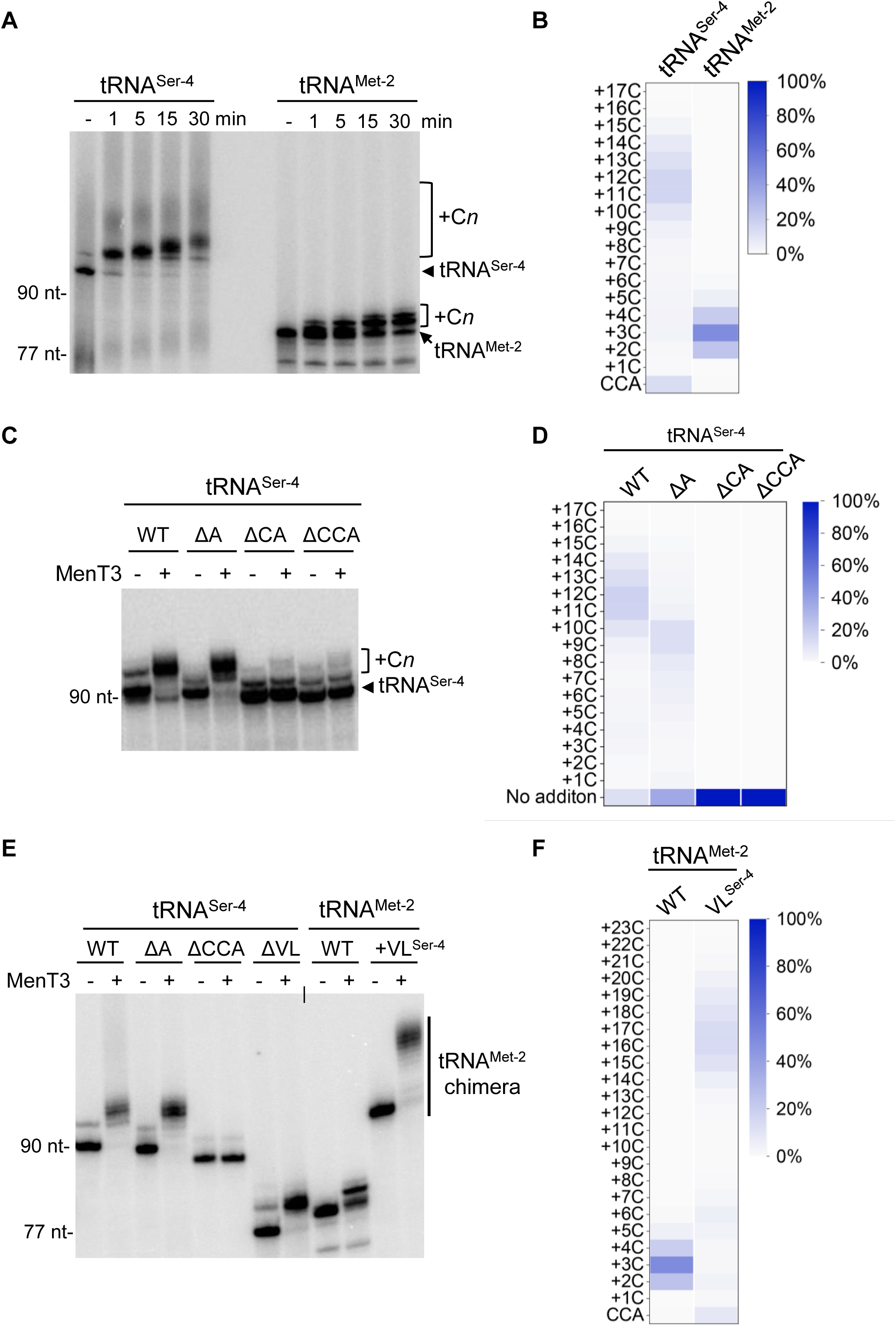
tRNA structural and sequence determinants for MenT3 modifications *in vitro.* (**A**) MenT3 tRNA preference *in vitro*. Purified α-P32 labeled tRNA^Ser-4^ and tRNA^Met-2^ were incubated in the presence of 1 mM CTP with MenT3 (0.2 µM) at 37 ℃ for different time points, separated on a 10% urea gel and revealed by autoradiography. (**B**) tRNA-Seq mapping of purified tRNA^Ser-4^ and tRNA^Met-2^. Purified tRNA^Ser-4^ and tRNA^Met-2^ (20 ng/µl) were incubated with MenT3 (1 µM) and 1 mM CTP for 10 min at 37 ℃ and subjected to tRNA-seq. (**C**) MenT3 modification of tRNA^Ser-4^ 3’-end length variants. Purified α-P32 labeled 3′ΔCCA, ΔCA, and ΔA truncated tRNA^Ser-4^ were incubated with MenT3 (0.2 µM) for 5 min at 37 °C, separated as in panel (A). (**D**) tRNA-seq mapping of truncated tRNA^Ser-4^ ΔCCA, ΔCA and ΔA variants from (B). (**E**) Impact of tRNA^Ser^ variable loop on MenT3 activity. Purified labeled tRNA^Ser-4^ deleted for its long variable loop (ΔVL) and the tRNA^Met-2^ ^(VLSer-4)^ chimera with the long variable loop of tRNA^Ser-4^ (+VL^Ser-4^) were incubated with MenT3 (0.2 µM) for 5 min at 37 °C, separated as in (A). (**F**) tRNA-Seq mapping of purified tRNA^Met-2^ and tRNA^Met-2(VLSer-4)^ chimera from panel (E) Purified tRNA (20 ng/µl) were incubated with MenT3 (1 µM) and 1 mM CTP for 10 min at 37 ℃ and subjected to tRNA-seq. Representative results of triplicate experiments are shown in in panel A, C and D, and sequencing data from two independent experiments are shown in panels in B, D and F (Duplicate is shown in datasheet file).

We next investigated the sequence and structural elements within a tRNA that are required for modification by MenT3 *in vitro*. Several variants of tRNA^Ser-4^ with sequential deletion or mutation of the 3′end nucleotides were labelled, purified using the ribozyme-based cleavage method and incubated with MenT3 in the presence of CTP. Under such conditions, we found that the adenosine amino acceptor of the CCA motif could be deleted or mutated to U, C, or G without detectably affecting MenT3 activity (**Fig. 2C and Supplementary Figure S1**). In sharp contrast, both 3′ΔCA and 3′ΔCCA end deletions within tRNA^Ser-4^ fully prevented modification by MenT3 (**Fig. 2C**). The 3′-OH tRNA-seq data confirmed that 3′ΔA, but not 3′ΔCA or 3′ΔCCA tRNA^Ser-4^, can be modified by MenT3 (**Fig. 2D**). Yet, we noticed that cytidine extensions on 3′ΔA were on average slightly shorter than for the wild type 3′CCA tRNA^Ser-4^ (*i.e.,* 9 to 10 cytidines compared 11 to 12), suggesting that the last nucleotide could still have some weak impact on MenT3 activity (**Fig. 2D**). Together these data demonstrate that 3′CCA and CCΔA ends are *bona fide* targets for MenT3.

The main structural feature that differentiates serine tRNAs from the other tRNAs in *M. tuberculosis* is the presence of a longer variable loop (Cai *et al*, 2020), which we believe could play a role in MenT3 specificity. To answer this, we first deleted the variable loop in tRNA^Ser-4^ and tested the resulting construct *in vitro* in the presence of MenT3. Remarkably, deletion of the variable loop of tRNA^Ser-4^ abolished the formation of long poly-C extensions, leaving the deleted form of the tRNA^Ser-4^ to be extended as per non-serine tRNAs of *M. tuberculosis* (**Fig. 2E**). We next asked whether grafting the variable loop of tRNA^Ser-4^ to an unrelated tRNA would induce the formation of long poly-C extensions by MenT3. We constructed a tRNA^Met-2^ chimera in which the native short variable loop was replace by the extended variable loop of tRNA^Ser-4^, and tested for extension in our *in vitro* assay. The data presented in **Fig. 2E** indeed show that the engineered tRNA^Met-2^ chimera with the tRNA^Ser-4^ variable loop is efficiently modified by MenT3 and accumulates long poly-C extensions, in a manner comparable to that of tRNA^Ser-4^ wild type. Analysis by 3′-OH tRNA seq confirmed that the tRNA^Met-2^ chimera contained long poly-C extensions (up to n=23) with a majority of 16 to 17 cytidines (**Fig. 2F**). Together these data demonstrate the importance of the variable loop of tRNA^Ser^ for MenT3 activity.

### tRNA^Ser^ is the target of MenT3 *in vivo* in *M. tuberculosis*

We next investigated the cellular tRNA targets of MenT3 *in vivo* in its native host *M. tuberculosis*. MenT3 was expressed in *M. tuberculosis* H37Rv Δ*menAT3* mutant strain for 0, 3 and 24 h at 37 °C, total RNAs were extracted and 3′-OH tRNA-seq was performed and compared to *M. tuberculosis* H37Rv wild type (**Fig. 3A**). Using this recently developed method (Xu *et al*, 2023), we were able to identify all of the 45 tRNAs of *M. tuberculosis* within the extracts (**Fig. 3B and Supplementary datasheet**). Strikingly, the analysis shows that MenT3 only targets tRNA^Ser^ isoacceptors *in vivo*. Modification of tRNA^Ser^ by 3′ addition of cytidines was very robust *in vivo* (approximately 65% of tRNA^Ser-2, 3, 4^ and 20% of tRNA^Ser-1^ as judged from three independent replicates) after 24 h of expression (**Supplementary Figure S2**). None of the other tRNAs were detectably modified by MenT3. Similar results were observed in *M. smegmatis* (**Supplementary Figure S3**). These data demonstrate that serine tRNAs are the targets of MenT3 in *M. tuberculosis.* Unexpectedly, analysis of the seq data revealed that different tRNA^Ser^ 3′-end species were accumulating *in vivo* than *in vitro* (**Fig. 3C**). First, we found that cytidine extensions were significantly shorter *in vivo*, with the majority of tRNA^Ser^ having only a single added cytidine and a small fraction reaching a maximum of n=5. In addition, while cytidine extensions occur after the adenosine nucleotide of the 3′CCA end (CCA+C*_n_*) *in vitro*, the vast majority, if not all, of the 3′ ends detected *in vivo* were deleted for A or CA, leading mainly to the accumulation of 3′CCΔA+C_(1-5)_ extensions but also to 3′CΔCA without added cytidine (on average 7% tRNA^Ser-4^, 27% tRNA^Ser-3^, 10% tRNA^Ser-2^ and 4% tRNA^Ser-1^ as judged from three independent replicates) after 24 h of expression, which corresponds to only one doubling time for *M. tuberculosis*. Note that the appearance of such species of tRNA^Ser^ *in vivo* was confirmed by northern blot using a probe against tRNA^Ser-3^ in *M. smegmatis* (**Supplementary Figure S3**). In addition, although not present in *M. tuberculosis*, the *in vivo* data confirm that tRNA^Sec^ can also be targeted by MenT3 in *M. smegmatis* (**Supplementary Figure S3 and Fig.1E**). Remarkably, the fact that 3′CΔCA ends are not modified by MenT3 *in vitro* (**Fig. 2C**) strongly suggests that such tRNA species could accumulate *in vivo* following the processing of MenT3-modified tRNAs by endogenous RNases counteracting a primary CCA+C*n* elongation of tRNA^Ser^ by MenT3 (see below). Accordingly, such RNases could also be responsible for generating 3′CCΔA ends that are efficiently processed by MenT3 *in vitro* (**Fig. 2C**) and which accumulate as the major 3′CCΔA+C_(1-5)_ species *in vivo* (**Fig. 3C**).

**Fig. 3:**
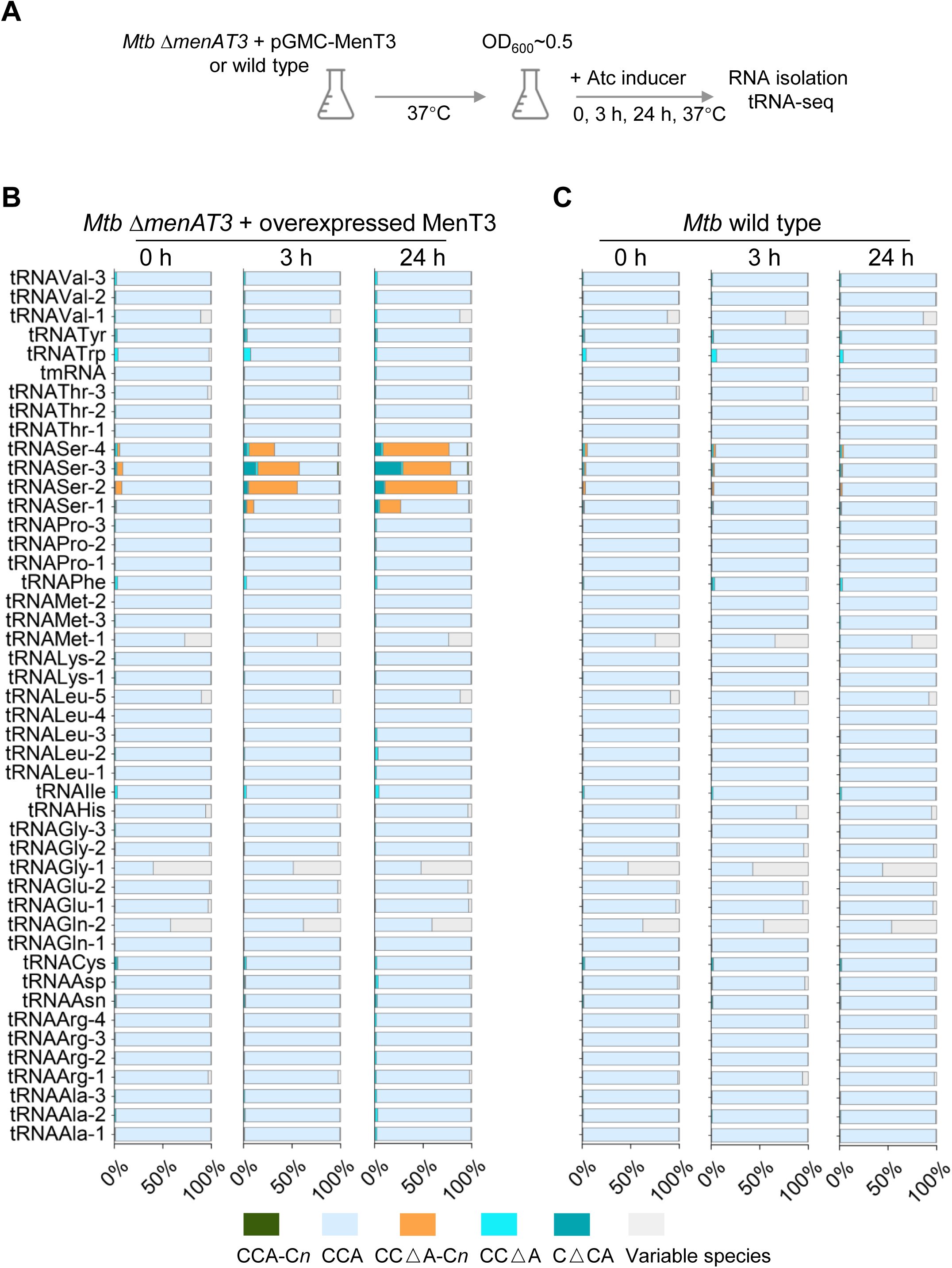
MenT3 specifically targets tRNA^Ser^ in *M. tuberculosis.* (**A**) Experimental conditions for tRNA-seq in *M. tuberculosis*. *M. tuberculosis* (*Mtb*) wild type H37Rv strain and its isogenic mutant *ΔmenAT3* expressing MenT3 from the integrative pGMC vector were individually grown at 37 °C in 7H9 medium supplemented with 10% albumin-dextrose-catalase (ADC, Difco) and 0.05% Tween 80. When the OD_600_ reached to about 0.5, the anhydrotetracycline inducer (Atc, 200 ng/ml) was added and cells were collected after 0, 3 or 24 h incubation at 37 °C. Total RNA was extracted and tRNA-seq was performed. The percentage of modified tRNA per tRNA species identified for the mutant overexpressing MenT3 (**B**) and the wild type strain (**C**) is shown. The names of the identified tRNA for both strains are shown on the left of panel B. The data are presented as the mean value obtained from three independent experiments. A detailed view of the different modification obtained for tRNA^Ser^ is shown in **Supplementary Figure S2**.

### *M. tuberculosis* RpH counteracts MenT3-mediated elongation of tRNA 3′-ends

In order to investigate the possible role of tRNA repair enzymes of *M. tuberculosis* that could generate 3′ CCΔA or CΔCA templates in response to MenT3, we tested the effect of the three main candidates present in *M. tuberculosis*, namely RpH, Orn and PNPase (Taverniti *et al*, 2011). The three ribonucleases were purified and individually incubated for 1 h with samples of labelled tRNA^Ser-4^ that have been previously modified by MenT3 and in which CTP has been removed to prevent possible residual NTase activity of MenT3 (**Fig. 4A**). Under such conditions, we found that the 3′ exonuclease RpH of *M. tuberculosis*, but not PNPase or Orn (**Fig. 4A**), was able to efficiently trim the poly-C modified tRNA^Ser-4^ to generate 3′ CCA, CCΔA and CΔCA ends (**Fig. 4B and 4C**). Indeed, while MenT3 alone could modify over 85% of the tRNA^Ser-4^ with CCA+C*_n_* ends, the addition of RpH led to the processing of most the MenT3-modified tRNA^Ser-4^ to generate wild type 3′-CCA (∼37%), CCΔA (∼36%) and CΔCA (∼11%,) ends (**Fig. 4B and 4C**). Furthermore, when the assay was performed with MenT3-modified tRNA^Ser-4^ 3′CCΔA mutant instead of the wild type tRNA^Ser-4^, the MenT3 generated 3′CCΔA+C*n* extensions could also efficiently be trimmed by RpH to produce 3′-CCΔA (60%) and CΔCA (30%) after RpH treatment **Fig. 5 D and E**). This shows that RpH can generate both 3′CCΔA ends that can be efficiently re-processed as substrate by MenT3 and 3′CΔCA ends, which are not further modified by MenT3 and thus would accumulate *in vivo* (**Fig. 3B**). Together these data suggest that RpH might be responding to the effect of the toxin by trimming the primary MenT3-modified 3′CCA+C*n* ends of tRNA^Ser^ in order to regenerate either 3′CCA or compatible ends for repair by CCA adding enzymes. Yet, the fact that 3′CCΔA is a *bona fide* MenT3 substrate would make such response ineffective in *M. tuberculosis* (see discussion).

**Fig. 4:**
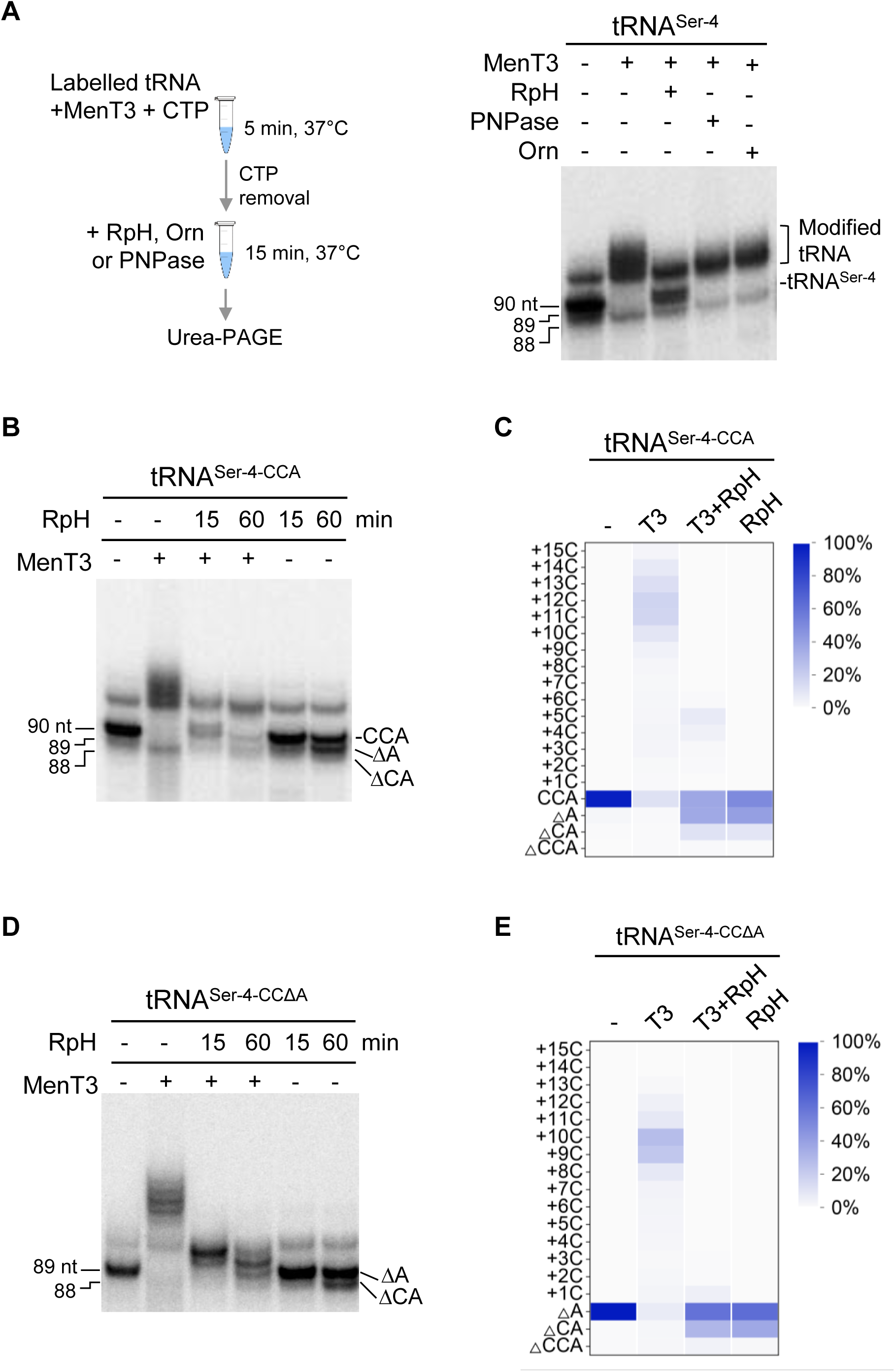
RpH-mediated response to MenT3 modifications *in vitro*. (**A**) tRNA repair assay used to test RpH, PNPase, and Orn of *M. tuberculosis*. Left panel was shown the developed tRNA repair assay; α-P32 labelled tRNA^Ser-4^ was incubated with MenT3 (0.2 µM) for 5 min at 37 ℃ and the modified tRNA was subjected to repair by RpH, PNPase, or Orn (10 µM) for 15 min at 37 ℃, the samples were separated on a 10% urea gel and revealed by autoradiography. (**B**) and (**C**) Repair of modified tRNA^Ser^ by RpH. The tRNA repair assay was performed as in (A), except that incubation was performed for 15 and 60 min at 37 ℃ in the presence and in the absence of MenT3. (**C**), tRNA-seq mapping of tRNA repaired by RpH. 20 ng/μl tRNA was incubated with MenT3 (1 μM) in the presence of 1 mM CTP for 10 min at 37 ℃. Following removal of CTP, the modified tRNA was then incubated with RpH (10 μM) for 1 h at 37 ℃ and tRNA-seq was performed. Similar *in vitro* reaction (**D**) and tRNA-seq experiment (**E**) were performed for tRNA^Ser-4-CCΔA^ as substrate. Representative results of triplicate experiments are shown in in panel A, B and D, and sequencing data from two independent experiments are shown in panels C and E (Duplicate are shown in datasheet file).

**Fig. 5:**
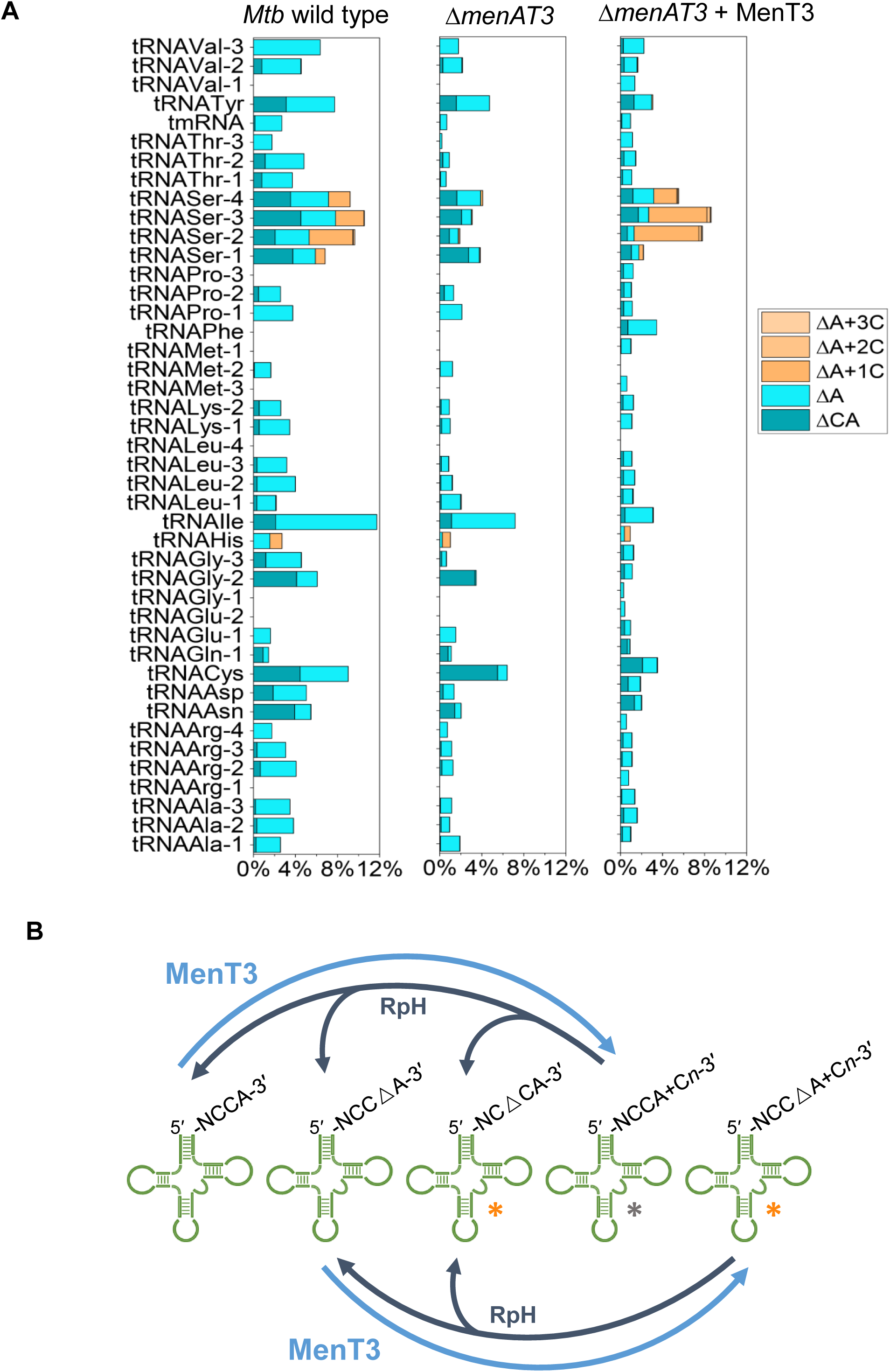
Steady state tRNA^ser^ modification in *M. tuberculosis* by endogenous MenT3. (**A**) *M. tuberculosis* wild-type H37Rv, the isogenic *ΔmenAT3* mutant and *ΔmenAT3-*menT3 complementation strain were individually grown at 37 °C in 7H9 medium supplemented with 10 % albumin-dextrose-catalase (ADC, Difco) and 0.05% Tween 80. When the OD_600_ reached to 0.5, the cells were collected. Total RNA was extracted to perform tRNA-seq. The percentage of modified tRNA per tRNA species identified is shown. The data are presented as the mean value obtained from three independent experiments. A detailed view of the different modification obtained for tRNA^Ser^ is shown in **Supplementary Figure S4**. (**B**) The model for MenT3 activity and its interplay with RpH in *M. tuberculosis* is described in the discussion section. The asterisks indicate tRNA products accumulating *in vivo* (orange) and *in vitro* (grey).

### Steady state tRNA^Ser^ modification in *M. tuberculosis*

The 3′-OH tRNA-seq analysis of the *M. tuberculosis* H37Rv wild type control under normal growth conditions (**Supplementary Figure S2**) revealed a significant steady state level of tRNA^Ser^ 3′-ends with MenT3-like modifications (about 8% of the total tRNA^Ser^). In this case, the only possible source of MenT3 would come from the native chromosomal *menAT3* operon (**Supplementary Figure S2)**. In order to investigate whether endogenous *menAT3* from wild type *M. tuberculosis* could indeed control the active pool of tRNA^Ser^ *in vivo*, we performed similar *in vivo* 3′-OH tRNA-seq experiments using the *M. tuberculosis* H37Rv wild type, its isogenic Δ*menAT3* mutant and the Δ*menAT3* mutant carrying MenT3 on the pGMC integrative plasmid in the absence of inducer, in order to obtain low level leaky expression of MenT3 and avoid toxicity. The data presented in **Fig. 5A** and **Supplementary Figure S4** show that indeed, the MenT3-like modifications of tRNA^Ser^ that are present in *M. tuberculosis* wild type decrease significantly in the Δ*menAT3* mutant. Note that MenT3 expression without inducer was sufficient to partly restore tRNA^Ser^ modifications in a Δ*menAT3* mutant background, yet less efficiently than in the presence of inducer (**Fig 5A compared to Fig 3B and 3C**). When comparing each of the tRNA^Ser^ isoacceptors (**Supplementary Figure S4**), the MenT3-like modifications in the Δ*menAT3* mutant dropped from 9.5 to 1.8% for tRNA^Ser-2^, from 10.5 to 3% for tRNA^Ser-3^, from 9.2 to 4.1% for tRNA^Ser-4^ and from 6.7 to 3.8% for tRNA^Ser-1^, which was also less efficiently modified *in vivo* following MenT3 overexpression (see **Fig. 3C**). Among these, the 3′CCΔA+C*n* ends of tRNA^Ser^ were barely detected in the Δ*menAT3* mutant (**Supplementary Figure S4**). Together these *in vivo* data indicate that there is a fraction of MenT3 toxin that is active under standard laboratory growth conditions that can modulate the pool of mature tRNA^Ser^ available for translation in wild type *M. tuberculosis*.

## DISCUSSION

This work demonstrates that the MenT3 toxin has a robust NTase activity *in vitro* and identifies its native tRNA targets *in vivo* in *M. tuberculosis*. It shows that MenT3 can modify all the tRNA tested *in vitro* by specifically adding cytidines to their 3′ end, although with a preference for tRNA^Ser^, to which significantly longer stretches of cytidines were added. Transcriptomic identification of MenT3 tRNA targets in *M. tuberculosis* revealed that tRNA^Ser^ isoacceptors are the sole targets of the toxin *in vivo* and that significantly different tRNA 3′ends accumulate *in vivo* and *in vitro*, most likely due to detoxification attempts by *M. tuberculosis* RpH in response to MenT3. Finally, this work identifies a basal level of MenT3-dependent tRNA^Ser^ 3′end modifications in *M. tuberculosis* under standard growth conditions, when MenT3 levels are physiological and produced from the native chromosomal *menAT3* operon.

The observation that MenT3 only targets tRNA^Ser^ isoacceptors *in vivo* and that cytidine elongations are significantly shorter *in vivo* (1 to 5) than *in vitro* (up to 17) suggests more stringent conditions *in vivo* or/and that other factors would contribute to such a strong preference *in vivo.* Remarkably, as observed in *E. coli* (Avcilar-Kucukgoze *et al*, 2016) and in humans (Evans *et al*, 2017), tRNA^Ser^ are significantly less aminoacylated than other tRNAs in *M. tuberculosis*, with less than 20% of tRNA^Ser^ being charged when at steady state (Tomasi *et al*, 2023). Such a low charging of tRNA^Ser^ was proposed to be due to a competition between aminoacylation of tRNA^Ser^ for translation and the need for serine amino acids for pyruvate and acetate production and glycine synthesis in *E. coli* (Avcilar-Kucukgoze *et al*, 2016). Therefore, the low availability of endogenous charged tRNA^Ser^ in *M. tuberculosis* would exacerbate the deleterious effect of MenT3 and contribute to the evolution of a strong preference for tRNA^Ser^ *in vivo*. The targeting of specific tRNAs is a characteristic of many toxin families (Songailiene *et al*, 2020; Yashiro *et al*, 2021; Tomasi *et al*, 2023; Cheverton *et al*, 2016; Wilcox *et al*, 2018; Zhang *et al*, 2020; Kurata *et al*, 2021; Li *et al*, 2021; Vang Nielsen *et al*, 2019). In *M. tuberculosis*, the acetyltransferase toxin TacT (Rv0919) was shown to specifically acetylate the primary amine group of charged tRNA^Gly^ glycyl-tRNAs (Tomasi *et al*, 2023), and the PIN domain RNase toxin VapC-mt4 was shown to cleave a single site within the anticodon sequence of tRNA^Cys^ (Barth *et al*, 2021). Collectively, these observations indicates that the selective targeting of a single tRNA or a small subset of tRNAs may be a hallmark of successful toxins of TA systems.

This work also revealed the importance of the long variable loop characteristic of tRNA^Ser^ for MenT3 activity (Throll *et al*, 2023). Noticeably, this region was shown to be important for binding to the Seryl-tRNA synthetase (SerRS) both in *E. coli* and in humans (Throll *et al*, 2023; Lenhard *et al*, 1999), suggesting that it could also contribute to the specific binding of MenT3.

We observed striking differences in the nature of the tRNA^Ser^ 3′end species that accumulated *in vivo* when compared to *in vitro*, *i.e*., mainly CCΔA+C*n* and CΔCA *in vivo* (with a small fraction of CCΔA, ΔCCA and CCA+C*n*), and exclusively CCA+C*n in vitro*. This strongly suggests that *in vivo* cytidine elongations are recognized and trimmed by *M. tuberculosis* tRNA repair enzymes, although not enough to restore a sufficient pool of mature tRNA. Accordingly, we have identified the 3′exoribonuclease RpH of *M. tuberculosis* as the main response to MenT3 activity. Although its role in tRNA maturation or repair in *M. tuberculosis* is unknown, RpH was shown to play an important role for the removal of nucleotides downstream of the 3′CCA end motif from tRNA precursors in other bacteria (Wen *et al*, 2005). In this case, processing by RpH not only led to the formation of compatible tRNA 3′CCA ends but also to a significant fraction of 3′CΔCA truncated tRNA ends (Wen *et al*, 2005), which is in line with the species accumulating *in vivo* and *in vitro* in the presence of *M. tuberculosis* RpH and MenT3. Therefore, together these data suggest a model (**Fig. 5B**) in which MenT3 first attacks uncharged, mature tRNA^Ser^ 3′CCA ends, adding cytidine extensions (CCA+C*n*) to block amino acylation. Such MenT3-modified elongated tRNAs represent *bona fide* clients for RpH, which can efficiently trim 3′ends to regenerate CCA, but also CCΔA and CΔCA ends. Since we showed that CΔCA is not a substrate of MenT3, such species may thus increasingly accumulate *in vivo* as a product of RpH attempts to repair elongated tRNA ends. Both CCA and CCΔA 3′ ends resulting from RpH activity will in turn be targeted by MenT3 to generate CCA+C*n* and CCΔA+C*n,* which can then be targeted again by RpH and enter a new cycle, progressively leading to a decrease in CCA+Cn species and the accumulation of CCΔA+C*n* and CΔCA species (**Fig. 5B**).

Previous data showed that *E. coli* is significantly less sensitive to *M. tuberculosis* MenT toxins (*i.e*., MenT1, 3, and 4) and that RpH overexpression can partially counteract MenT3 toxicity in *E. coli*, but not in *M. smegmatis* (Cai *et al*, 2020). Although we cannot exclude that *E. coli* and *M. tuberculosis* RpH respond differently to cytidine addition, a reasonable hypothesis for such differences might be that most of the tRNA genes of *M. tuberculosis* (30/45) do not contain a CCA motif in their sequences and need to be further processed by the essential CCA-adding enzyme (Błaszczyk *et al*, 2023). This is in sharp contrast with *E. coli*, in which the CCA-adding enzyme is not essential and all the tRNA genes have the CCA in their sequence (Zhu & Deutscher, 1987). As a result, upon expression of MenT3 the CCA-adding enzyme of *M. tuberculosis* might be overwhelmed by the new pool of accumulating tRNA^Ser^ CCΔA+C*n* and CΔCA 3′-ends.

Our data highlight the existence of a fraction of tRNA^Ser^ that is modified by MenT3 *in vivo* in *M. tuberculosis* wild type under standard growth conditions. This unexpected discovery suggests the existence of a population of endogenous MenT3 that is active and that could modulate translation, as part of a significant contribution to the control of pathogen growth during infection.

In agreement with this hypothesis, a screen for transposon mutants that failed to grow in murine macrophages identified *menT3*, and not *menA3*, as one of the 126 genes important for *M. tuberculosis* intracellular survival under such conditions (Rengarajan *et al*, 2005). More recently, *menT3* was also identified as a persistence gene in *M. tuberculosis*, which contributes to the long term survival in mouse lungs (Dutta *et al*, 2014), thus further supporting the requirement of endogenous MenT3 activity under relevant physiological conditions for this pathogen.

The fact that MenT3 can be active and target tRNA^Ser^ despite the presence of the MenA3 antitoxin is intriguing and might be related to its peculiar mode of inhibition. Indeed, previous work showed that MenA3 acts as a type VII antitoxin kinase, which transiently interacts with, and inhibits, MenT3 by phosphorylating the MenT3 S78 catalytic site residue (Yu *et al*, 2020; Cai *et al*, 2020). Accordingly, this inhibitory mechanism was recently supported by phospho-proteome analysis of *M. tuberculosis*, which indeed identified the presence of phosphorylated MenT3 at S78 (Frando *et al*, 2023). Therefore, one attractive possibility is that a fraction of inactive phosphorylated MenT3 could be dephosphorylated in response to certain growth conditions or aggression by the host immune system. In support of this, the housekeeping phosphatase PstP of 1. *M. tuberculosis* could dephosphorylate MenT3 *in vitro* (Yu *et al*, 2020). Reversibly, a fraction of free active MenT3 could also be affected by endogenous Ser/Thr protein kinases (STPKs) like PknD and PknF (Frando *et al*, 2023), suggesting that the level of active endogenous MenT3 toxin might not solely be controlled by the antitoxin, but also by responsive endogenous kinase/phosphatase networks to ensure translational control is attuned to physiological need during growth and infection.

## MATERIALS AND METHODS

### Bacterial strains

*E. coli* strains DH5α (Invitrogen), BL21(λDE3) and BL21 (λDE3) AI (Novagen), *M tuberculosis* H37Rv (WT; ATCC 27294) and its mutant derivative H37Rv △*menAT3::dif6/pGMCZ* and *M. smegmatis* mc^2^155 (ATCC 700084) have been described (Cai *et al*, 2020). *E. coli* were grown at 37 °C in LB, when necessary, with kanamycin (Km, 50 μg.ml^-1^), ampicillin (Ap, 50 μg.ml^-1^), isopropyl-β-D-thiogalactopyranoside (IPTG, 1 mM), L-arabinose (L-ara, 0.1 % w/v) or D-glucose (glu, 0.2 % w/v). *M. smegmatis* mc^2^155 was grown at 37 °C in LB, when necessary, with Km (10 μg.ml^-1^) or streptomycin (Sm, 25 μg.ml^-1^). *M. tuberculosis* strains were grown at 37 °C in 7H9 medium (Middlebrook 7H9 medium, Difco) supplemented with 10 % albumin-dextrose-catalase (ADC, Difco) and 0.05 % Tween 80 (Sigma-Aldrich), or on complete 7H11 solid medium (Middlebrook 7H11 agar, Difco) supplemented with 10 % oleic acid-albumin-dextrose-catalase (OADC, Difco). When necessary, media were supplemented with, hygromycin (Hy, 50 μg.ml^-1^), Sm (25 μg.ml^-1^), zeocin (Zeo, 25 μg.ml^-1^), or anhydrotetracycline (Atc, 100 or 200 ng.ml^-1^) (Xu *et al*, 2023).

### Plasmid constructs

Plasmids pET15b (Novagen), pET20b and pETDuet-1 have been described. All the primers used to construct the plasmids are described in **Supplementary Table S1**. To construct pET15b-menT3, *rv1045* was PCR amplified from the *M. tuberculosis* H37Rv genome using primers Rv1045 NdeI-For and Rv1045 BamHI-Rev and cloned into pET15b after digestion with NdeI and BamHI enzymes. For pET20b-mtbrpH or pET20b-mtborn construction, *rv1340* or *rv2511* was PCR amplified from the *M. tuberculosis* H37Rv genome using primers Rv1340 NdeI-For and Rv1340 XhoI-Rev or Rv2511 NdeI-For and Rv2511 XhoI-Rev, respectively. Amplified fragments were then inserted into pET20b following digestion with NdeI and XhoI enzymes. To construct pETDuet-1-mtbpnpase, *rv2783c* was PCR amplified from the *M. tuberculosis* H37Rv genome using primers Rv2783c MfeI-For and Rv2783c HindIII-Rev, and cloned as MfeI/ HindIII fragments into EcoRI/HindIII digested pETDuet-1.

The pUC57-T7-Met-2-HDV and pUC57-T7-Ser-4-HDV plasmids containing tRNA-HDV fusion under the control of a T7 promoter were synthesized by Genewiz (Azenta Life Sciences). The pUC57-T7-Met-2+variable loop of Ser-4-HDV, pUC57-T7-Ser-4ΔA-HDV, pUC57-T7-Ser-4ΔCA-HDV, pUC57-T7-Ser-4ΔCCA-HDV and pUC57-T7-Ser-4Δvariable loop-HDV plasmids were constructed by PCR using the primers listed in **Supplementary Table S1**. The sequences of the T7-Met-2-HDV, T7-Met-2+variable loop of Ser-4-HDV, T7-Ser-4-HDV and T7-Ser-4Δvariable loop-HDV fragments are given in **Supplementary Table S1**. To prepare DNA template for transcription with T7 polymerase, T7-tRNA-HDV fragments were PCR amplified using primers T7-For and HDV-Rev for T7-Met-2-HDV or HDV short-Rev for T7-Met-2+variable loop of Ser-4-HDV, T7-Ser-4-HDV and T7-Ser-4Δvariable loop-HDV. T7-Ser-4ΔA-HDV, T7-Ser-4ΔCA-HDV and T7-Ser-4ΔCCA-HDV was constructed by In-fusion PCR and the primers were listed in **Supplementary Table S1**.

### Protein expression and purification

Purified MenT3 was produced as described previously (Cai *et al*, 2020). Purified proteins were also obtained as follows; to purify MenT3, mtbRpH, mtbPNPase and mtbOrn, strain BL21(λDE3) AI transformed with pET15b-MenT3, pET20b-RpH, pETDuet-1-RV2783c or pET20b-Orn was grown to an OD_600_ of approximately 0.4 at 37 °C, 0.2 % L-ara was added and the culture immediately incubated overnight at 22 °C, respectively. Cultures were centrifuged at 5000 x *g* for 10 min at 4 °C, pellets were resuspended in lysis buffer (50 mM Tris-HCl, pH8.0, 200 mM NaCl, 10 mM MgCl_2_, 20 mM imidazole; 20 ml for 1 liter of cell culture) and incubated for 30 min on ice. Lysis was performed using the One-shot cell disrupter at 1.5 Kbar (One shot model, Constant Systems Ltd). Lysates were centrifuged for 30 min at 30000 x *g* in 4 °C and the resulting supernatants were gently mixed with Ni-NTA Agarose beads (Qiagen) pre-equilibrated with lysis buffer, at 4 °C for 30 min in a 10 ml poly-prep column (Bio-Rad). The column was then stabilized for 10 min at 4 °C, washed five times with 10 ml of lysis buffer, and proteins were eluted with elution buffer (50 mM Tris-HCl, pH8.0, 200 mM NaCl, 10 mM MgCl_2_, 250 mM imidazole). 500 µl elution were collected and PD MiniTrap G-25 columns (GE Healthcare) were used to exchange buffer (50 mM Tris-HCl, pH8.0, 200 mM NaCl, 10 mM MgCl_2_, 10 % glycerol) and proteins were concentrated using vivaspin 6 columns with a 5000 Da cut off (Sartorius). To remove the His-tag, thrombin was incubated with the protein at 4 °C overnight. Following NTA and streptavidin addition, the cleaved His-tag and thrombin were washed out. Proteins were stored at –80 °C until further use.

### *In vitro* transcription of tRNAs

tRNAs were synthesized via *in vitro* transcription using PCR templates that incorporated an integrated T7 RNA polymerase promoter sequence. Primers for *M. tuberculosis* tRNAs are given in **Supplementary Table S1**. The T7 RNA polymerase *in vitro* transcription reactions were carried out in a total volume of 25 µl, which included a 5 µl nucleotide mix containing 2.5 mM NTPs (Promega). For each reaction, 50 ng to 100 ng of template DNA were used, along with 1.5 µl of rRNasin (40 U.ml-1, Promega), 5 µl of 5x optimized transcription buffer (Promega), 2 µl of T7 RNA polymerase (20 U.ml^-1^), and 2.5 µl of 100 mM DTT. The reactions were incubated at 37 °C for 2 hours (Cai *et al*, 2020). The resulting tRNA products were extracted using Trizol reagent(Yip *et al*, 2020) and stored at a final concentration of 100 to 200 ng.µl^-1^, as determined by NanoDrop analysis. To obtain high-concentration tRNA for EMSA, the TranscriptAid T7 High Yield Transcription Kit from Thermo Fisher was employed. In this protocol, 1 µg of tRNA DNA template was included in the transcription assay and incubated at 37 °C for 4 hours. Subsequently, DNase I treatment was performed, followed by RNA isolation using Trizol.

### *In vitro* transcription of tRNAs with homogeneous 3′ ends

An optimized version of the hepatitis delta virus (HDV) ribozyme was used to generate homogeneous tRNA 3′ ends as described (Schürer *et al*, 2002; Xu *et al*, 2023). Briefly, the DNA template T7-tRNA-HDV was amplified from plasmid pUC-57Kan-T7-tRNA-HDV (Supplementary Table S1). Labelled or unlabelled tRNAs were prepared by *in vitro* transcription of PCR templates using T7 RNA polymerase. The T7 RNA polymerase *in vitro* transcription reactions were performed in 25 µl total volume, with a 5 µl nucleotide mix of 2.5 mM ATP, 2.5 mM UTP, 2.5 mM GTP, 60 µM CTP (Promega, 10 mM stock) and 2-4 µl 10 mCi.ml^-1^ of radiolabelled CTP [α-32P], or with 5 µl nucleotide mix of 2.5 mM ATP, 2.5 mM UTP, 2.5 mM GTP, 2.5 mM CTP for unlabelled tRNA transcription. 50 to 100 ng of template were used per reaction with 1.5 µl rRNasin 40 U.ml^-1^ (Promega), 5 µl 5x optimized transcription buffer (Promega), 2 µl T7 RNA polymerase (20 U.ml^-1^) and 2.5 µl 100 mM DTT. Unincorporated nucleotides were removed by Micro Bio-Spin 6 columns (Bio-Rad) according to manufacturer’s instructions. The transcripts were gel-purified on a denaturing 6 % acrylamide gel and eluted in 0.3 M sodium acetate overnight at 20 °C. The supernatant was removed, ethanol precipitated and resuspended in 14 µl nuclease-free water. Radioactively labelled tRNAs carrying a 2′,3′ cyclic phosphate at the 3′ end was dephosphorylated using T4 polynucleotide kinase (NEB) in 100 mM Tris-HCl pH 6.5, 100 mM magnesium acetate and 5 mM β-ME in a final volume of 20 µl for 6 h at 37 °C. All assays were desalted by Micro Bio-Spin 6 columns (Bio-Rad).

### Electrophoretic mobility shift assay (EMSA)

The concentrations of proteins are indicated in the figures and figure legends, and tRNA was consistently maintained at a concentration of 50 ng. µl^-1^ in all experimental setups. The tRNA– MenT3 complexes were assembled in a buffer solution comprising 20 mM Tris-HCl (pH 8.0), 10 mM NaCl, 5% glycerol, and 10 mM MgCl_2_. The reactions underwent incubation at room temperature for a duration of 30 min and were subsequently subjected to resolution through Tris-Borate-EDTA (TBE) native Polyacrylamide Gel Electrophoresis (PAGE).

### Nucleotide transfer assay

MenT3 NTase activity was assayed in 10 µl reaction volumes containing 20 mM Tris-HCl pH 8.0, 10 mM NaCl, 10 mM MgCl_2_ and 1 µCi.µl^-1^ of radiolabelled rNTPs [α-^32^P] (Hartmann Analytic) and incubated for 5 min at 37 °C. 100 ng *in vitro* transcribed tRNA product, 1 µg total RNA or 100 ng of *E. coli* or *M. smegmatis* tRNA was used per assay with 0.2 µM of protein. The 10 µl reactions were purified with Bio-Spin® 6 Columns (Bio-Rad), and mixed with 10 µl of RNA loading dye (95% formamide, 1 mM EDTA, 0.025 % SDS, xylene cyanol and bromophenol blue), denatured at 90 °C and separated on 6 % polyacrylamide-urea gels. The gel was vacuum dried at 80 °C, exposed to a phosphorimager screen and revealed by autoradiography using a Typhoon phosphorimager (GE Healthcare).

For the single tRNA modification by MenT3 *in vitro*, the radiolabeled tRNA was incubated with 0.2 µM MenT3 in the presence of 1 mM CTP for 5 min at 37 °C, the reaction was halted by adding 10 µl of RNA loading dye, denatured at 90 °C. Subsequently, the modified RNA samples were separated on a 10% polyacrylamide-urea gel.

### tRNA repair *in vitro*

The tRNA modified product from MenT3 *in vitro* reactions underwent the following steps: it was initially purified using Bio-Spin® 6 Columns (Bio-Rad), followed by incubation with a putative exoribonuclease (10 µM) in a reaction buffer (For Orn: 10 mM Tris-HCl, pH 8.0, 100 mM NaCl, 5 mM MgCl_2_, For RpH and PNPase: 50 mM Tris-HCl, pH 8.0, 10 mM NaCl, 2.5 mM MgCl_2_, 10 mM K_2_HPO_4_, 1 mM DTT) at 37 °C for an indicative time. Subsequently, it was mixed with 10 µl of RNA loading dye, subjected to denaturation at 90 °C, and then separated using 10% polyacrylamide-urea gels. Alternatively, the RNA samples were analyzed through RNA-seq.

### Northern blot

Total RNA was isolated from *M. smegmatis* transformed with either pGMC-MenT3 or an empty vector. Subsequently, 5 µg of RNA was separated using 10 % polyacrylamide-urea gels. Following gel electrophoresis, the RNA was transferred onto nylon membranes (Amersham Hybond N+, GE Healthcare) and subjected to hybridization as previously described(Fricker *et al*, 2019). The sequences of the oligonucleotides used are described in **Supplementary Table S1.**

### tRNA libraries and sequencing

Primers used for the construction of tRNA libraries are described in **Supplementary Table S1**. To obtain the MenT3 library from *in vitro* reactions, 5 µg of *M. smegmatis* total RNA supplemented with 1 mM CTP was incubated with water and 0.8 µg MenT3, then incubated for 20 min at 37 °C. For the *in vitro* transcribed tRNA-seq, 20 ng.µL^-1^ of specific tRNAs were incubated with 1 µM MenT3 at 37 °C for 10 min. Total RNA samples and single tRNA samples were isolated using trizol and ethanol precipitation, respectively. Construction of tRNA-seq libraries from *M. smegmatis* and *M. tuberculosis* were performed as follows: *M. smegmatis* transformed with pGMC or pGMC-menT3 weak RBS were grown at 37 °C for 3 days, then the culture was transferred to fresh LB, until OD_600_ reached 0.1, Atc (100 ng.ml^-1^) was added, and the cells were collected after 3 hours at 37 °C for preparation of total RNA, as previously described (Xu *et al*, 2023). *M. tuberculosis* wild-type H37Rv, H37Rv *ΔmenAT3* mutant or the same strains transformed with pGMC-menT3 weak RBS were grown at 37 °C until OD_600_ reached 0.5, Atc (200 ng.ml^-1^) was added and the cells were collected at 0, 3 or 24 h. Cell pellets were resuspended in 1ml of Trizol and cells were disrupted in a bead-beater disrupter, after addition of glass beads. Samples were centrifuged for 2 min. at 20,000 g and the trizol extract was collected and conserved for at least 48 hours at –80°C before transfer out of the BSL3 laboratory for total RNA isolation. All of the RNA was dissolved in DEPC-H_2_O (pH 7.0) by heat at 65 °C for 10 min, which conditions known to be able to discharge the tRNA *in vitro*(Walker & Fredrick*, 2008). To remove m1A, m3C and m1G modifications in the tRNAs, the total RNA samples were pretreated using the demethylase kit (Arraystar, cat#: AS-FS-004, Rockville, MD, USA), followed by trizol isolation. Notably, during the demethylation reaction (pH 7.5-8)(Toh *et al*, 2020), tRNA was deacylated sufficiently by this near-neutral pH. The 3p-v4 oligo was 5′ adenylated using 5′ DNA Adenylation Kit (E2610S, NEB) according to the manufacturer’s protocol. To construct the library, RNA samples were ligated to the adenylated 3p-v4 adaptors, and reverse transcription was performed with ProtoScript II RT enzyme (NEB) using barcode primers (**Supplementary Table S1**). Finally, PCR amplification was performed with tRNAs oligoFor mix and A-PE-PCR10 (after 5 cycles, the program was paused and B_i7RPI1_CGTGAT or i7RPI7_ GATCTG was added) using Q5 Polymerase Hot-Start (NEB). The library was sequenced by DNBSEQ –G400RS High-throughput Sequencing Set (PE150) in BGI Genomics (Hong Kong).

### RNA-Seq data processing

After a quality check, reads were demultiplexed to obtain a fastq file per experimental condition. For each experimental condition, the same procedure was applied: (i) mapping to a reference, (ii) PCR duplicates removal, (iii) quantification of read counts. Raw reads quality was checked with FastQC (http://www.bioinformatics.babraham.ac.uk/projects/fastqc). Further reads processing was performed with R software version 4.3.1 and BioConductor libraries for processing sequencing data obtained. R library ShortRead 1.58.0(Morgan *et al*, 2009) was used to process fastq files. Rsubread 2.14.2(Liao *et al*, 2019) was used to map reads on the reference genome. Further filtering was performed to ensure reads contained the structure resulting from library preparation (**Supplementary Table S1** for RNA-Seq read structure): reads start with a random nucleotide at position 1, a valid barcode at position 2-5, a recognition sequence resulting from ligation at position 6-24 with one mismatch allowed, random UMI sequence at position 25-39, agacat control sequence at position 40-45, and a random nucleotides sequence at position 46-150, which corresponds to the reverse complement of the ligated 3′-end of tRNAs in the experiment. Reads corresponding to the different experimental conditions identified by the barcode were demultiplexed into fastq files for independent further analyses (mapping and quantification).

All the *M. smegmatis* mc^2^155 (NC_008596.1) and *M. tuberculosis* H37Rv (NC_000962.3) chromosomal tRNA gene sequences were extracted in a multifasta file. The sequence of genes *rrfA*, cspA and for tmRNA were also added. The nucleotides CCA were concatenated at the end of each sequence. Mapping on reference sequences was performed by using the library Rsubread 2.14.2 with parameters ensuring that the read maps without any indel to a unique region: unique=TRUE, type=’dna’, maxMismatches=2, indels=0, ouput_format=’BAM’.

For quantification, SAM files were processed with Rsamtools 2.16.0 (https://bioconductor.org/packages/Rsamtools) to count the number of reads mapping to the same region in reference sequences with the same 3′ unmapped sequences due to RNA 3′ end modifications. Reads were kept if the aligned region nearest the 3’ end of the reference sequence was at least 20 nucleotides long and allowing 3’ end soft clipping (unaligned region after the minimum 20 nucleotides aligned nearest the 3’ end of the reference sequence), which corresponds to nucleotides added by the toxin. In the alignments in the SAM file, this translates to a CIGAR code ending with only ‘M’ characters (indicating a match) possibly followed by ‘S’ characters (indicating no alignment to the reference sequence). PCR duplicates were removed using the UMI introduced in the library preparation for reads mapping exactly the same region (same beginning and end of the alignment) and having exactly the same unmapped 3′-end sequence. After these preprocessing steps, reads were regrouped by their 3′-end mapping position and unmapped 3′-end region sequence to quantify their abundance in terms of same tRNA 3′-end followed by the same nucleotides *i.e.* same post-transcriptional modifications. To remove noise, a tRNA with its post-transcriptional modifications was kept only if it was found at least 10 times in at least one of the experimental conditions.

Primers and specific sequences used in this work are shown in **Supplementary Table S1**

## DATA AVAILABILITY

All the data needed to evaluate the conclusions in the paper are present in the paper and/or in the Supplementary Datasheet.

## Supporting information

supplementary datasheet

## ACKNOWLEDGEMENTS

We thank Ciarán Condon for useful discussion. This work was supported by the Centre National de la Recherche Scientifique, Université Paul Sabatier, the Programme d’Investissements d’Avenir (ANR-20-PAMR-0005) to ON and PG; the Swiss National Science Foundation (CRSII3_160703) to PG; the National Natural Science Foundation of China, (32000021) to XX; a scholarship from the China Scholarship Council (CSC) as part of a joint international PhD program with –Toulouse University Paul Sabatier and a Fondation pour la Recherche Médicale (FDT202304016729) to X.H.; a Springboard Award (SBF002\1104) from the Academy of Medical Sciences to BU and TRB; an Engineering and Physical Sciences Research Council Molecular Sciences for Medicine Centre for Doctoral Training studentship (EP/S022791/1) to T.J.A.

## AUTHOR CONTRIBUTIONS

– Analyzed data: X.X, R.B, B.V, T.J.A, B.U, C.G, X.H, P.R, T.R.B, O.N., P.G.

– Designed research: X.X, R.B, B.V, T.J.A, B.U, C.G, X.H, P.R, T.R.B, O.N., P.G.

– Performed research: X.X., R.B, B.V, T.J.A, B.U, C.G, X.H

– Wrote the paper: X.X and P.G. with contributions from all the authors.

– Funding acquisition: X.X, O.N., T.R.B. and P.G.

– Supervised the study: P.G.

