## Supplementary file for "Nucleotidyltransferase toxin MenT targets and extends the aminoacyl acceptor ends of serine tRNAs *in vivo* to control *Mycobacterium tuberculosis* growth"

This file contains 4 Supplementary Figures and 1 Table

A

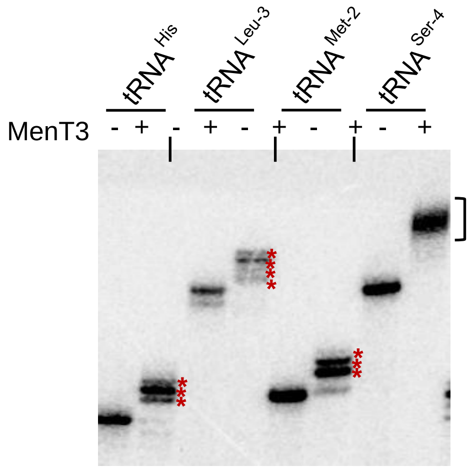


B

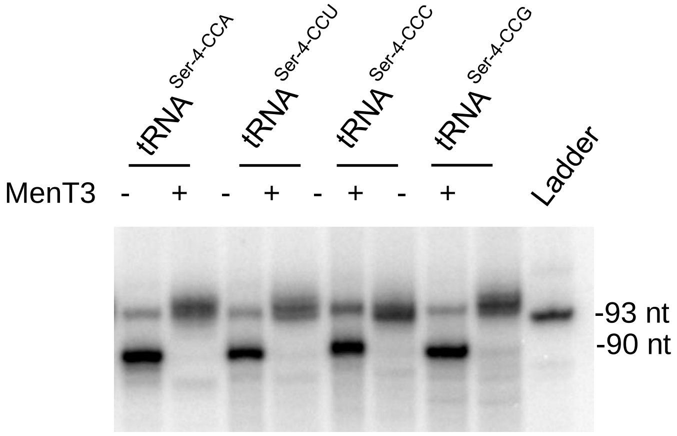


**Supplementary Figure S1:** (**A**) Data extended from Figure 2 (A). Purified α-P32 labeled tRNA$^{Ser-4}$ and tRNA$^{Met-2}$, tRNA$^{His}$ and tRNA$^{Leu-3}$ were incubated in the presence of 1 mM CTP with MenT3 (0.2 µM) at 37 °C for 5 min, separated on a 10% urea gel and revealed by autoradiography. (**B**) Purified α-P32 labeled tRNA$^{Ser-4}$ with mutations in the 3′ adenosine of the CCA end were incubated in the presence of 1 mM CTP with MenT3 (0.2 µM) at 37 °C for 5 min, separated on 10% urea gel and revealed by autoradiography. Representative results of triplicate experiments are shown.

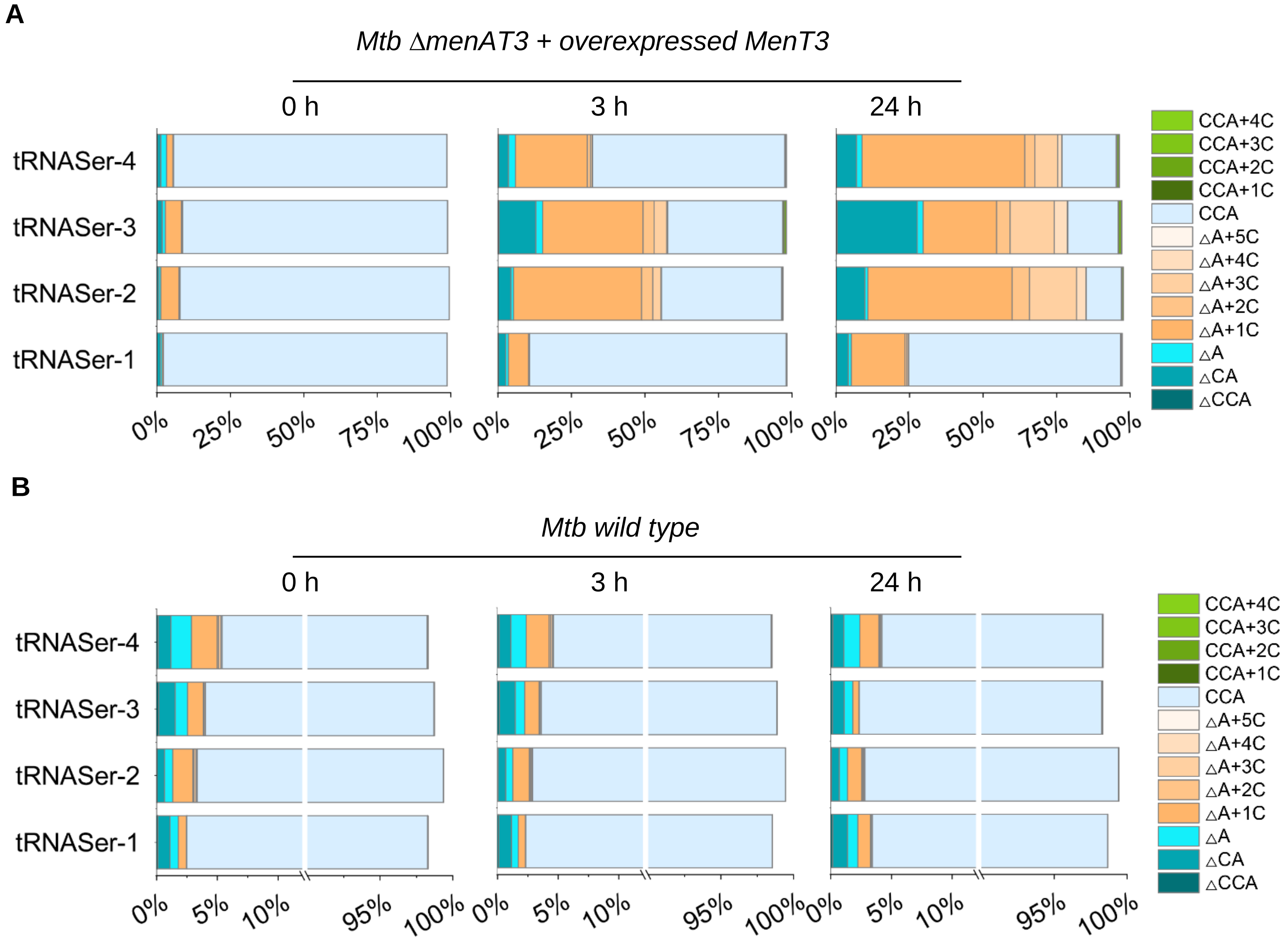


**Supplementary Figure S2:** A detailed analysis of the tRNA seq data of the 3′-end of tRNA$^{\text{Ser-1, 2, 3 and 4}}$ of *M. tuberculosis* wild-type H37Rv strain and its isogenic mutant Δ*menAT3* expressing MenT3. The data were mean analyzed from three independent experiments.

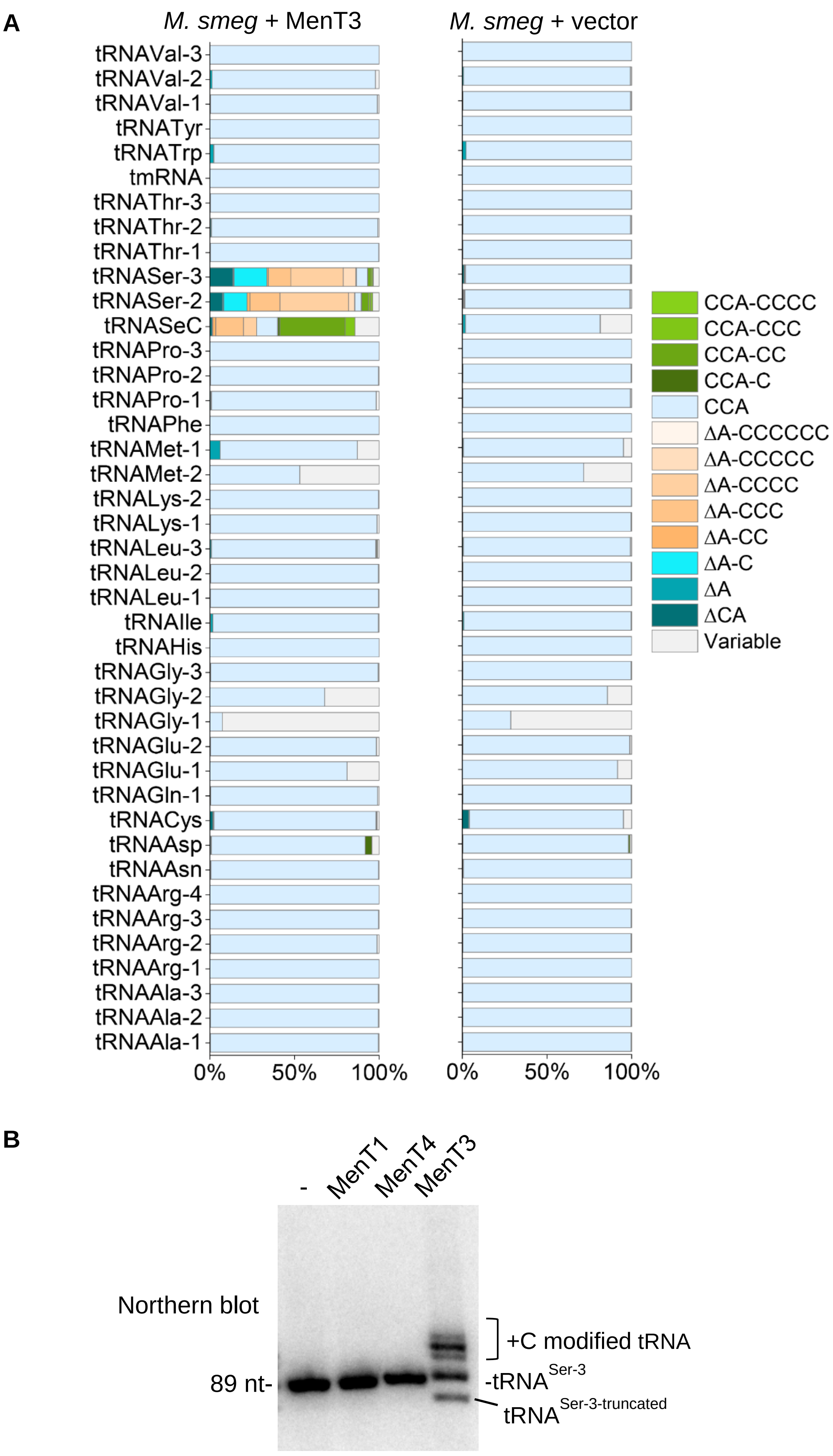
A
M. smeg + MenT3
M. smeg + vector
tRNAVal-3
tRNAVal-2
tRNAVal-1
tRNATyr
tRNATrp
tmRNA
tRNAThr-3
tRNAThr-2
tRNAThr-1
tRNASer-3
tRNASer-2
tRNASeC
tRNAPro-3
tRNAPro-2
tRNAPro-1
tRNAPhe
tRNAMet-1
tRNAMet-2
tRNALys-2
tRNALys-1
tRNALeu-3
tRNALeu-2
tRNALeu-1
tRNAIle
tRNAHis
tRNAGly-3
tRNAGly-2
tRNAGly-1
tRNAGlu-2
tRNAGlu-1
tRNAGln-1
tRNACys
tRNAAsp
tRNAAsn
tRNAArg-4
tRNAArg-3
tRNAArg-2
tRNAArg-1
tRNAAla-3
tRNAAla-2
tRNAAla-1
0%
50%
100%
CCA-CCCC
CCA-CCC
CCA-CC
CCA-C
CCA
ΔA-CCCCCC
ΔA-CCCCC
ΔA-CCCC
ΔA-CCC
ΔA-CC
ΔA-C
ΔA
ΔCA
Variable
B
-
MenT1
MenT4
MenT3
Northern blot
89 nt-
+C modified tRNA
-tRNA Ser-3
tRNA Ser-3-truncated

**Supplementary Figure S3:** (**A**) tRNA-seq analysis in *M. smegmatis* overexpressing MenT3. *M. smegmatis* containing empty vector or pGMC-menT3 were individually grown at 37 °C in LB supplemented with 0.05% Tween 80. When the $OD_{600}$ reached to 0.1, the anhydrotetracycline inducer (Atc, 100 ng/ml) was added and cells were collected after 3 h incubation at 37 °C. Total RNA was extracted and tRNA-seq was performed as in **Fig 3**. Results are from two independent experiments (Duplicate is shown in datasheet file). (**B**) The *M. smegmatis* RNA extracts from (A) were subjected to northern blot analysis using a probe against $tRNA^{Ser\text{-}3}$ (labelled by α-P32) and revealed by autoradiography. The results revealed the presence of elongated $tRNA^{Ser\text{-}3}$ as well as a truncated $tRNA^{Ser}$ was observed on a TBE-Urea gel, likely corresponding to the $tRNA^{Ser}$-CΔCA identified in the tRNA-seq data. Representative results of duplicate experiments are shown.

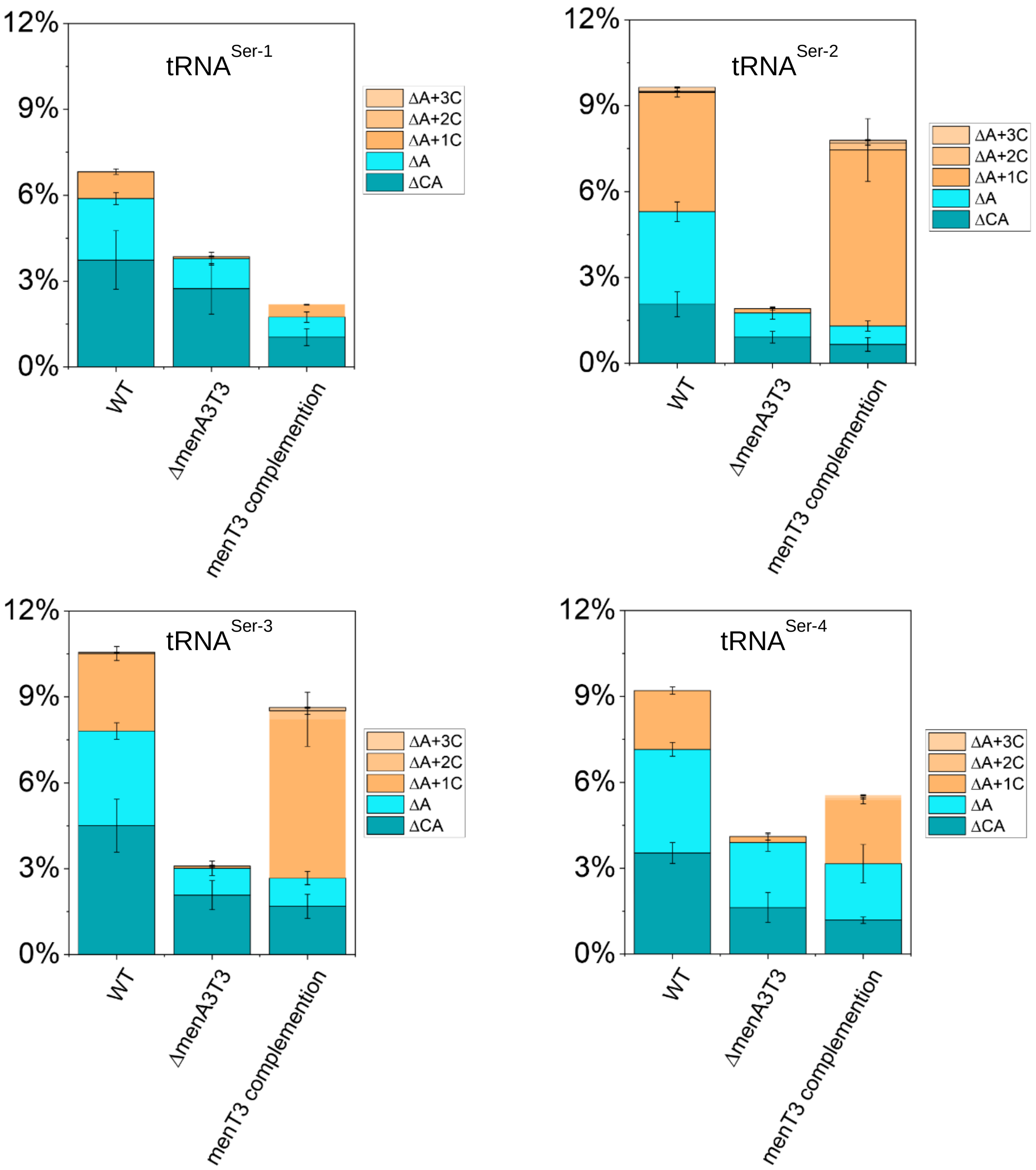


**Supplementary Figure S4.** Details of the modifications found for four tRNA$^{Ser}$ isoacceptors were analyzed among wild-type *M. tuberculosis*, Δ*MenAT3* and menT3 complementation strain. The data shown as means of three independent experiments.

**Supplementary Table S1:** Primers and specific sequences used in this work

| **Gene name** | **Primers** | **Primer sequence (5′ to 3′)** |
|---|---|---|
| pET15b-menT3 | Rv1045 NdeI-For | ttcatatgaccaagccctattcgtcgc |
| | Rv1045 BamHI-Rev | ttggatcctcatcttttcgtcgcccga |
| pET20b-mtbrpH | Rv1340 NdeI-For | ttcatatgtccaagcgagaagacggc |
| | Rv1340 XhoI-Rev | ttctcgagggtgccaaacgccttcggc |
| pET20b-mtborn | Rv2511 NdeI-For | ttcatatgcaagacgaactggtgtgga |
| | Rv2511 XhoI-Rev | ttctcgagaccgctctggggcgcctcg |
| pETDuet-1-mtbpnpase | Rv2783c MfeI-For | ttcaattggtctgccgctgaaattga |
| | Rv2783c HindIII-Rev | ttaagctttcagctggtgaccgtcgcg |
| pUC57-T7-Met-2+variable loop of Ser-4-HDV | Met2insertvariloop-Ser4F | gacggctaacaccgtctcgccggttcgagtccgg |
| | Met2insertvariloop-Ser4R | acggtgttagccgtctaacgattatgagtcgttcgc |
| pUC57-T7-Ser-4Δvariable loop-HDV | Ser4 variloop delFor | agcgagagccgggggttcaaatccctcc |
| | Ser4 variloop deleRev | cccccggctctcgctttcaaggcgagtgc |
| **tRNA primers** | | |
| Mtb tRNA His | Mtb tRNA His For | attaatacgactcactataggtgagtgtagttcagttggt |
| | Mtb tRNA His Rev | tggggtgagtgacgggactc |
| Mtb tRNA Leu-3 | Mtb tRNA Leu-3 For | attaatacgactcactataggtccgagtggcggaatggcagacgcgcta |
| | Mtb tRNA Leu-3 Rev | tggtgtccgaggggggactt |
| Mtb tRNA Met-2 | Mtb tRNA Met-2 For | attaatacgactcactatagggcgatgtagctcagtcggttagagcga |
| | Mtb tRNA Met-2 Rev | tggtagcgatggccggactcgaa |
| Mtb tRNA Ser-4 | Mtb tRNA Ser-4 For | attaatacgactcactatagggtggcgtggcagagcggcctaat |
| | Mtb tRNA Ser-1/4 Rev | tggcggtggcggagggattt |
| T7-tRNA | T7-For | attaatacgactcactat |
| | HDV-Rev | aaacgacggccagtgccaag |
| | HDV short-Rev | cttctcccttagcctaccg |
| pUC57-T7-Met-2+variable loop of Ser-4-HDV | Met2insertvariloop-Ser4F | gacggctaacaccgtctcgccggttcgagtccgg |
| | Met2insertvariloop-Ser4R | acggtgttagccgtctaacgattatgagtcgttcgc |
| | Ser4 variloop ΔFor | agcgagagccgggggttcaaatccctcc |

| | | |
|---|---|---|
| pUC57-T7-Ser-4Δvariable loop-HDV | Ser4 variloop ΔRev | cccccggctctcgctttcaaggcgagtgc |
| pUC57-T7-Ser-4ΔA-HDV | Ser4CCAΔA For | ccaccgccgggtcggcatggcatctcc |
| | Ser4CCAΔA Rev | ccgacccggcggtggcggagggatttg |
| pUC57-T7-Ser-4ΔCA-HDV | Ser4CCAΔCA For | gccaccgcgggtcggcatggcatctc |
| | Ser4CCAΔCA Rev | ccgacccgcggtggcggagggatttg |
| pUC57-T7-Ser-4ΔCCA-HDV | Ser4ΔCCA For | cgccaccggggtcggcatggcatctc |
| | Ser4ΔCCA Rev | ccgaccccggtggcggagggatttg |
| Northern blot | Ser-4-probe | ttcaaggcgagtgcattaggcc |
| | Ser-3-probe | ttccaggcgagcgccataggcc |
| | Gly-3-probe | cgcgggttcgattcccgtc |
| **Sequences of T7-tRNA-HDV constructs** | | |
| T7-Met-2-HDV<br>attaatacgactcactataggcgatgtagctcagtcggttagagcgaacgactcataatcgttaggtcgccggttcgagtccggccatcgctaccagggtcggcatggcatctccacctcctcgcggtccgacctgggctacttcggtaggctaagggagaagcttggcactggccgtcgttt | | |
| T7-Met-2+variable loop of Ser-4-HDV<br>attaatacgactcactataggcgatgtagctcagtcggttagagcgaacgactcataatcgttagacggctaacaccgtctcgccggttcgagtccggccatcgctaccagggtcggcatggcatctccacctcctcgcggtccgacctgggctacttcggtaggctaagggagaag | | |
| T7-Ser-4-HDV<br>attaatacgactcactataggtggcgtggcagagcggcctaatgcactcgccttgaaagcgagagacggctaacaccgtccgggggttcaaatccctccgccaccgccagggtcggcatggcatctccacctcctcgcggtccgacctgggctacttcggtaggctaagggagaag | | |
| T7-Ser-4Δvariable loop-HDV<br>attaatacgactcactataggtggcgtggcagagcggcctaatgcactcgccttgaaagcgagagccgggggttcaaatccctccgccaccgccagggtcggcatggcatctccacctcctcgcggtccgacctgggctacttcggtaggctaagggagaag | | |
| **RNA Seq primers** | | |
| Adaptor | 3p-v4 | Phosphate-***gtatct***<u>*nnnnnnnnnnnnnnTGAG*</u>cctcggttggtgccg-/SpacerC3/ |
| Barcode | D6A | ctctttccctacacgacgctcttccgatctntacacggcaccaaccgagg |
| | D6B | ctctttccctacacgacgctcttccgatctngtatcggcaccaaccgagg |
| | D6C | ctctttccctacacgacgctcttccgatctncgtccggcaccaaccgagg |
| | D6D | ctctttccctacacgacgctcttccgatctnaagtcggcaccaaccgagg |
| | D6E | ctctttccctacacgacgctcttccgatctnacaccggcaccaaccgagg |
| | D6F | ctctttccctacacgacgctcttccgatctnggtacggcaccaaccgagg |
| | D6H | ctctttccctacacgacgctcttccgatctntcggcggcaccaaccgagg |
| | D6I | ctctttccctacacgacgctcttccgatctncaagcggcaccaaccgagg |
| | D6J | ctctttccctacacgacgctcttccgatctnttgacggcaccaaccgagg |
| | D6K | ctctttccctacacgacgctcttccgatctngctgcggcaccaaccgagg |
| | D6L | ctctttccctacacgacgctcttccgatctnccgacggcaccaaccgagg |
| | D6M | ctctttccctacacgacgctcttccgatctnctcgcggcaccaaccgagg |
| | D6N | ctctttccctacacgacgctcttccgatctnaggacggcaccaaccgagg |
| | D6O | ctctttccctacacgacgctcttccgatctnattgcggcaccaaccgagg |
| | D6P | ctctttccctacacgacgctcttccgatctngacgcggcaccaaccgagg |
| | D6Q | ctctttccctacacgacgctcttccgatctntgttcggcaccaaccgagg |

| | | |
|---|---|---|
| | D6R | ctctttccctacacgacgctcttccgatctngcaacggcaccaaccgagg |
| | D6S | ctctttccctacacgacgctcttccgatctntcgtcggcaccaaccgagg |
| PCR | EMOTE-B_i7RPI7_GATCTG | caagcagaagacggcatacgagatgatctggtgactggagttcagacgtgtgc |
| | EMOTE-B_i7RPI1_CGTGAT | caagcagaagacggcatacgagatcgtgatgtgactggagttcagacgtgtgc |
| | **tRNAs oligoFor mix** | |
| | Ala tRNA seqFw | ctggagttcagacgtgtgctcttccgatct ggggctatggcgcagttggta |
| | Asn- Asp- Phe-Trp tRNA seqFw | ctggagttcagacgtgtgctcttccgatct gtagctcaattggcaagagc |
| | Cys tRNA seqFw | ctggagttcagacgtgtgctcttccgatct gtggccgagtggtgaggca |
| | Gln tRNA seqFw | ctggagttcagacgtgtgctcttccgatctaattggcaacacagctgatt |
| | Glu tRNA seqFw | ctggagttcagacgtgtgctcttccgatctcccccatcgtctagtggcct |
| | LeuCAG GAG TAG tRNA seqFw | ctggagttcagacgtgtgctcttccgatctgtggcggaatggcagacgcg |
| | LeuCAA TAA tRNA seqFw | ctggagttcagacgtgtgctcttccgatctcgtatcccaactggcagagg |
| | Lys tRNA seqFw | ctggagttcagacgtgtgctcttccgatctgttagctcagttggtagagc |
| | Met tRNA seqFw | ctggagttcagacgtgtgctcttccgatctgttagatcggcgggctcata |
| | Pro tRNA seqFw | ctggagttcagacgtgtgctcttccgatctcgggctgtggcgcagcttgg |
| | Sec tRNA seqFw | ctggagttcagacgtgtgctcttccgatctggaggcgtatccggtctggt |
| | SerGGA CGA TGA tRNA seqFw | ctggagttcagacgtgtgctcttccgatctagtggcctaaggcgctcgcct |
| | SerGCT Tyr tRNA seqFw | ctggagttcagacgtgtgctcttccgatctggaggcttgcccgagcggcc |
| | Thr tRNA seqFw | ctggagttcagacgtgtgctcttccgatctcttagctcagtcggcagagc |
| | Val tRNA seqFw | ctggagttcagacgtgtgctcttccgatctcgagtagctcagcgggagagc |
| | Arg tRNA seqFw | ctggagttcagacgtgtgctcttccgatctgcccccgtagctcaggggat |
| | Gly His tRNAs seqFw | ctggagttcagacgtgtgctcttccgatctgcgggcgtagctcaatggta |
| | Ile tRNAs seqFw | ctggagttcagacgtgtgctcttccgatctgggcctatagctcaggcggt |
| | Met-2 tRNA seqFw | ctggagttcagacgtgtgctcttccgatct ggcgatgtagctcagtcggt |
| | Ser-1/4 tRNA seqFw | ctggagttcagacgtgtgctcttccgatctggtggcgtggcagagcggcc |
| | tmRNA seqFw | ctggagttcagacgtgtgctcttccgatctttgagggaatgccgtagga |
| | 5s RNA seqFw | ctggagttcagacgtgtgctcttccgatct agctaagcctgccagcgccg |
| | A-PE-PCR10 | aatgatacggcgaccaccgagatctacactctttccctacacgacg |
| **RNA-Seq read structure** | | |

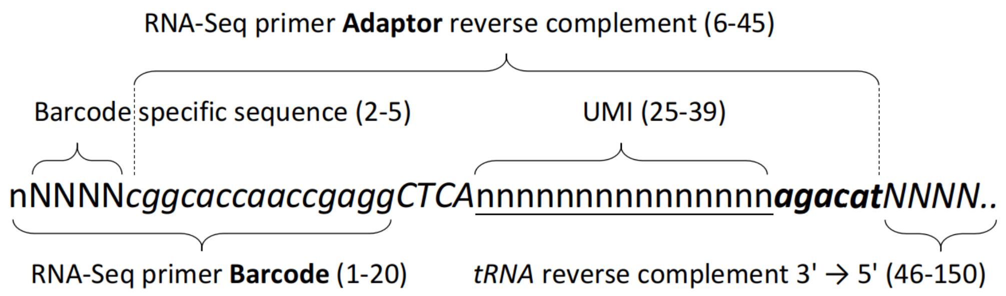


Position 1: random nucleotide

Position 2-5: RNA-Seq Barcode specific sequence from primer D6E, D6F, D6O, ..., in capital in the Barcode sequences

Position 6-24: sequence resulting from RNA-Seq Adaptor ligation i.e., end of Barcode (in italic in the Barcode sequence) followed by CTAC, the reverse complement of Adaptor (TGAG in capital in the Adaptor sequence)

Position 25-39: Unique Molecule Identifier (UMI) for PCR duplicate removal, underlined in the Adaptor sequence

Position 40-45: end of Adaptor reverse complement, agacat, in the RNA-Seq read (gtatct beginning of Adaptor in bold)

Position 46-150: reverse complement of the tRNA 3'end sequence
